# Soil microbes influence the ecology and evolution of plant plasticity

**DOI:** 10.1101/2024.10.23.619911

**Authors:** Lana G. Bolin

## Abstract

- Stress often induces plant trait plasticity, and microbial communities also alter plant traits. Therefore, it is unclear how much plasticity results from direct plant responses to stress versus indirect responses due to stress-induced changes to soil microbial communities.
- To test how microbes and microbial community responses to stress affect the ecology and potentially the evolution of plant plasticity, I grew plants in four stress environments (salt, herbicide, herbivory, no stress) with microbes that had responded to these same environments or with sterile inoculant.
- Plants delayed flowering under stress only in the presence of live microbial communities, and this plasticity was maladaptive. However, microbial communities responded to stress in ways that accelerated flowering across all environments. Microbes also affected the expression of genetic variation for plant flowering time and specific leaf area, as well as genetic variation for plasticity of both traits, and disrupted a positive genetic correlation for plasticity in response to herbicide and herbivory stress, suggesting that microbes may affect the pace of plant evolution.
- Together, these results highlight an important role for soil microbes in plant plastic responses to stress and suggest that microbes may alter the evolution of plant plasticity.

## INTRODUCTION

Stress (i.e., a condition that decreases a plant’s fitness below its potential; Osmond *et al*., 1987) is increasing with global change, and plant stress tolerance is often greatly enhanced, or in some cases made possible at all, by microbes (Rodriguez & Redman, 2008). For example, symbiotic microbes such as nitrogen-fixing bacteria, arbuscular mycorrhizal fungi, and endophytic fungi can buffer plants from nitrogen, phosphorus, and heat stress, respectively (Rodriguez *et al*., 2008; Smith & Read, 2010), and whole diverse soil and foliar microbial communities also appear to commonly increase plant stress tolerance (e.g., Xi *et al*., 2018; Fitzpatrick *et al*., 2019; Beals *et al*., 2020). Furthermore, microbial communities can respond rapidly to stress through shifts in community composition or through the evolution of key taxa (Elena & Lenski, 2003; Graves *et al*., 2015), and these microbial responses can benefit plant hosts under stress in phenomena known as “microbe-mediated adaptation” (sensu Petipas *et al*., 2021), “microbial rescue” (sensu Mueller *et al*., 2020), or “habitat-adapted symbiosis” (sensu Rodriguez *et al*., 2008). For example, rhizosphere microbes from saline environments improve plant salt stress tolerance (Yuan *et al*., 2019), soil fungi from serpentine environments promote plant growth and nutrient uptake in these nutrient-limited soils (Doubková *et al*., 2012), and microbial communities appear to commonly respond to drought in ways that increase plant fitness in dry environments (Lau & Lennon, 2012; Fitzpatrick *et al*., 2018; Giauque *et al*., 2019; Allsup & Lankau, 2019; Bolin *et al*., 2022; Ricks & Yannarell, 2023). On the other hand, microbial responses to stress could be antagonistic to plant responses if pathogens become more abundant under stress, or if stress reduces the effectiveness of plant defense against these pathogens (reviewed in Pandey & Kumar, 2019).

Microbes can affect plant stress tolerance via several mechanisms (reviewed in Kumar & Verma, 2018; Petipas *et al*., 2021; Angulo *et al*., 2022), one of which is by promoting phenotypic plasticity (hereafter, plasticity). Plasticity, or the production of alternate phenotypes in response to differing environments (DeWitt & Scheiner, 2004), is common in plants. When plant phenotypes shift in response to stress, we often assume that plants are responding to the stress environment directly, via what I call “direct plasticity” (Fig. **1a**). However, plant plasticity can also be mediated by microbes (Friesen *et al*., 2011; Goh *et al*., 2013). Therefore, if microbes rapidly respond to stress and alter plant phenotypes, microbes could be cryptically driving (or hindering) observed plant plasticity in response to stress. Such microbe-mediated plasticity can occur in two ways (Fig. **1b,c**). First, microbes that have responded to stress could cause plant traits to shift independently of the contemporary environment (e.g., Friesen *et al*., 2011; Jin *et al*., 2012; Lau & Lennon, 2012; Wagner *et al*., 2014; Panke-Buisse *et al*., 2015; Brunner *et al*., 2015) (Fig. **1b**). In this case the microbial community is the true environmental factor determining plant phenotype, such that plants produce alternate phenotypes in response to the composition of the soil microbial community regardless of the prevailing environmental conditions (and the stress environment itself plays no role). Alternatively, plants may only express plasticity (or express stronger plasticity) in response to stress in the presence of microbial communities that have responded to that stressor; without such a microbial community, plants would not express plasticity (or would express weaker plasticity) (Fig. **1c**; comparing red and black points) (reviewed in Goh *et al*., 2013). This could occur if plants rely on microbes they encounter under stress to cue their plastic response (Metcalf *et al*., 2019), or if these microbes produce chemicals that mimic plant hormones that induce plasticity (Friesen *et al*., 2011).

**Figure 1.**
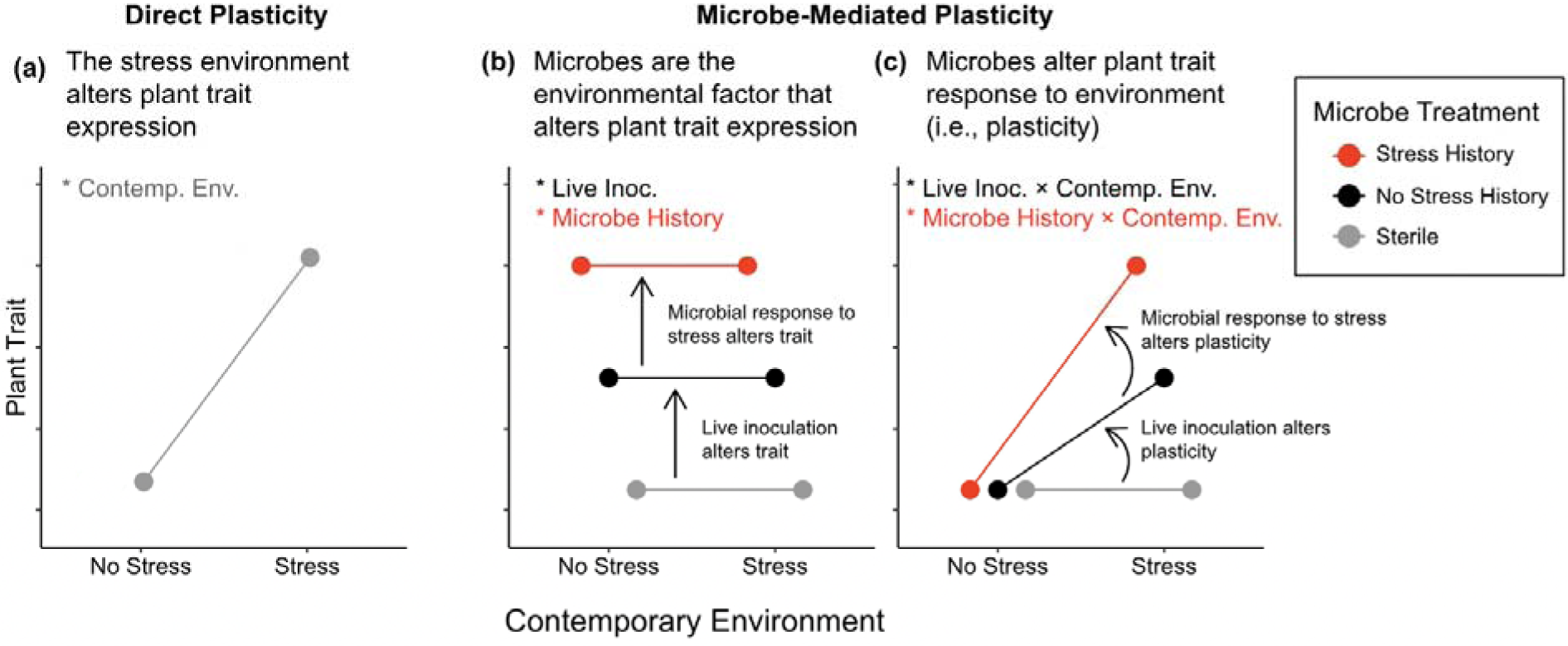
Graphical theory depicting (a) direct plasticity and (b, c) microbe-mediated plasticity. Plant ecologists would typically test for plasticity by measuring plant trait expression in two environments and attributing any change in the trait to the contemporary stress environment (a). However, microbes can rapidly respond to new environments, and may be mediating the observed plasticity in one of two ways: (b) microbes may be the true environmental factor that is altering plant trait expression, while the contemporary environment has no effect at all, or (c) microbes may alter plant trait responses to the environment (i.e., plasticity) such that the contemporary environment and the microbial community interact to determine plant phenotype. Hypothetical plants inoculated with microbes that have responded to stress are shown in red, plants inoculated with microbes from a non-stressful environment are shown in black, and plants inoculated with sterile inoculant are shown in gray. As shown, both live inoculation (comparing black to gray) and microbial responses to stress (“microbe history”; comparing red to black) mediate plasticity, but it could be the case that live inoculation mediates plasticity while microbe history does not, and vice versa. Statistical evidence for each type of plasticity is indicated at the top of each panel (asterisk indicates the term is significant; gray, black, and red text show hypothetical results from models comparing contemporary environments, live vs. sterile microbial treatments, and microbe history treatments, respectively).

Microbe-mediated plasticity of either type may buffer plants from the negative fitness effects of stress if they induce adaptive plant responses (i.e., if plant traits shift in the direction favored by natural selection), or could exacerbate the fitness effects of stress if plastic responses are maladaptive (i.e., if trait shifts are opposite the direction favored by natural selection) (Jordan, 1991; Nagy, 1997). In addition to the particular microbial communities encountered under stress, plant plasticity may be mediated by soil microbes generally such that their plasticity is dampened in a sterile environment (Fig. **1b,c**; comparing grey and black points).

In addition to affecting the expression of plasticity, microbes may also contribute to the evolution of plasticity in plants. Just like any other trait, the evolution of plasticity requires genetic variation, and will be facilitated or constrained by genetic correlations with other traits under selection. Microbes may affect both processes. First, there are conflicting hypotheses about how unfavorable conditions will affect the expression of genetic variation, with predictions ranging from greater, to lesser, to equal variation under stress relative to favorable conditions (Hoffmann & Merilä, 1999). Therefore, if a particular microbial community alters how favorable an environment is for plants, this could lead to changes in the expression of genetic variation in plant populations. If microbes ultimately increase the expression of plant genetic variation under stress, then microbes could reveal cryptic genetic variation that is “hidden” under conditions normally experienced by plants (e.g., Rutherford, 2003; Ghalambor *et al*., 2007; Ledón-Rettig *et al*., 2010). Although this new variation is likely to be mostly deleterious (e.g., Rutherford & Lindquist, 1998; Queitsch *et al*., 2002; Ghalambor *et al*., 2007), adaptive evolution may occur if some genotypes exhibit beneficial trait values or plasticity in the presence of microbes or particular microbial communities (Rutherford, 2003; Ghalambor *et al*., 2007). Second, just as there are ecological constraints to the *expression* of plasticity in response to multiple stressors (Valladares *et al*., 2007), there can also be genetic constraints to the *evolution* of plasticity in response to multiple stressors if plasticity in response to one stress is genetically correlated with plasticity in response to a different stress. Microbes may affect the evolution of plant plasticity by disrupting these genetic correlations if they alter plant traits in environment-specific ways (e.g., Goh *et al*., 2013), promoting divergent trait responses to different environments. Conversely, microbes may reinforce genetic correlations if their effects on plant traits are uniform across environments. Strong positive genetic correlations could slow evolution in response to multiple stressors if selection favors divergent phenotypes in different stress environments, or could accelerate the evolution of plasticity if selection is uniform across stressors.

Here, I investigated the effects of soil microbes on the ecology and evolution of plant plasticity under stress. I exposed replicated plant populations to three stressors and the soil microbial communities those stressors select for to test how microbial community responses to stress (microbe history) and the presence of diverse microbial communities (live vs. sterile inoculation) influence the effects of stress on plant plasticity. I then tested whether this plasticity (whether direct or microbe-mediated) was adaptive by comparing my findings to previous work that quantified the strength and direction of natural selection in this system. Finally, I tested whether microbial communities have the potential to alter the evolution of plant traits and plasticity by influencing (1) the expression of genetic variation for plant traits and trait plasticity, and (2) genetic correlations for plasticity across stress environments.

## MATERIALS AND METHODS

### Experimental Overview

To test how stress, live microbial inoculation, and microbial community responses to stress (microbe history) influence plant trait expression, the expression of genetic variation for traits and trait plasticity, and genetic correlations for plasticity, I collected live soil inoculant from a field experiment that included four stress treatments: salt, herbicide, simulated herbivory, and an unstressed control (“microbe history” hereafter; n = 3 randomized field plots per stress treatment; Fig. 2). I then planted *Chamaecrista fasciculata* (*Fabaceae*) into mesocosms inoculated with these field soils in the greenhouse and applied the same four stress treatments in a full factorial design (“contemporary environment” hereafter). I also included a sterile inoculant treatment, which allowed me to estimate the general effect of live field microbes (apart from their stress history) on plant traits.

**Figure 2.**
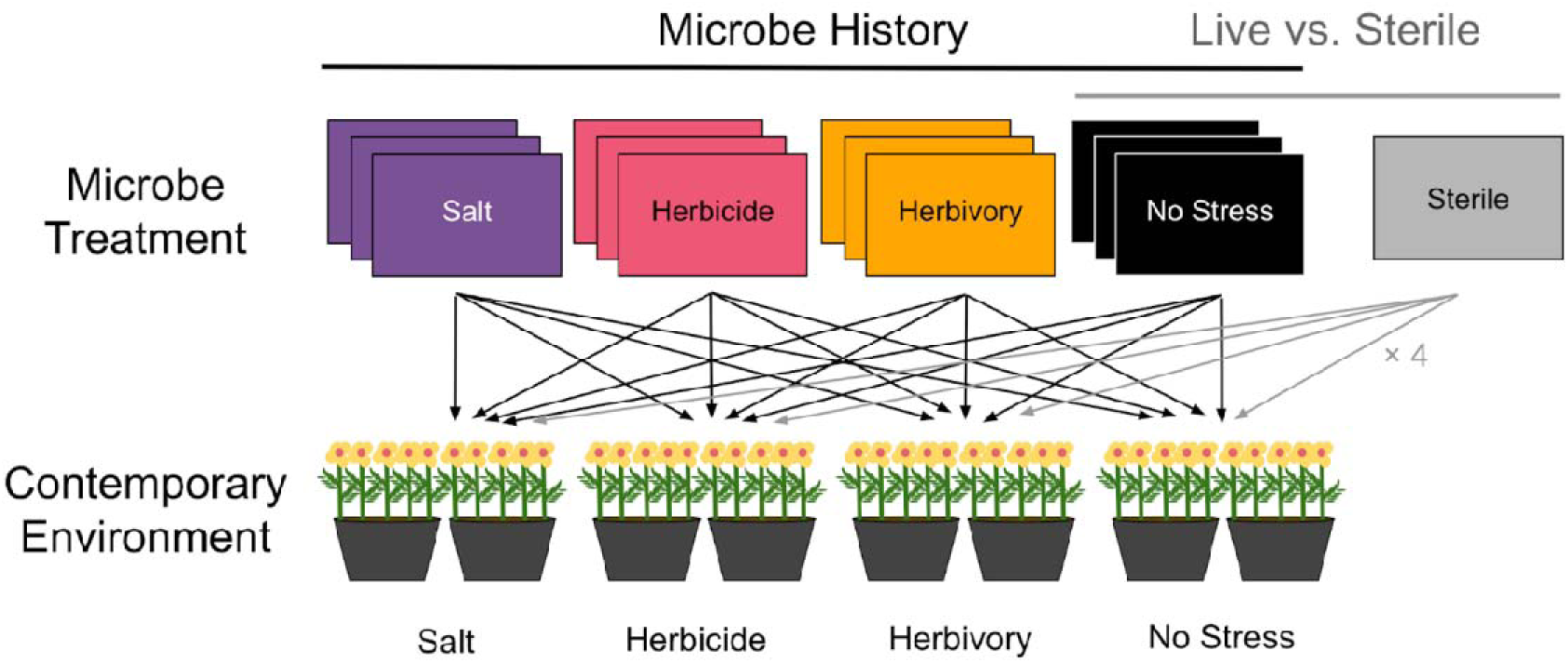
Experimental design. Triplicates of rectangles represents randomized field plots that received one of four stress treatments (“Microbe History”): salt stress (purple), herbicide stress (pink), simulated herbivory stress (gold), or no stress (black). The gray rectangle represents a sterilized combination of these field soils. Soil from each field plot was inoculated onto eight mesocosms that were treated with these same four stressors (“Contemporary Environment”) in a full factorial design, and the sterile inoculum was applied to 32 mesocosms (n = 8 per contemporary environment). An individual from each of 50 full-sibling families was randomly assigned to a location in each microbe × contemporary environment treatment; these 50 families were spread across two mesocosms, with 25 plants per mesocosm. N = 3,200 plants; n = 128 mesocosms. Live vs. sterile comparisons compare sterile inoculum to inoculum from the “no stress” field plots. Figure redrawn from Bolin & Lau (2024).

### Field-sourced microbial inoculants

I collected soil microbial inoculants from a field experiment that took place at the W.K. Kellogg Biological Station (KBS, Hickory Corners, Michigan, USA) in the summer of 2018. The field experiment was comprised of *C. fasciculata* populations (100 plants per 2 × 2 m field plot) planted into randomized field plots that were treated with 4 stress treatments: salt, herbicide, simulated herbivory, or an unstressed control (n = 3 field plots per stress treatment). For salt-stressed plots, 50 mL of 12 g/L, 24 g/L, and 24 g/L NaCl concentration was applied to the base of each plant 23, 38, and 45 d after planting, respectively. For herbicide-stressed plots, plants were sprayed twice with glyphosate (0.02 kg active ingredient/acre applied 29 d after planting, and 0.03 kg active ingredient/acre applied 40 d after planting). For simulated herbivory-stressed plots, all fully expanded leaves were removed with scissors 23-27 days after planting and again 38 days after planting. Note that the herbivory stress treatment was applied to the plant leaves so any changes to microbial communities were entirely mediated by the plant, while the salt and herbicide stress treatments include both direct effects on microbial communities as well as plant-mediated indirect effects. At the end of the growing season, I collected 3 liters of soil from the rhizosphere of *C. fasciculata* plants harvested from each field plot by shaking soil off the roots to use as inoculant in my greenhouse experiment.

### Greenhouse experiment

I sterilized 5-gallon pots (Zarn Inc 2000x, Reidsville, NC, USA) in 0.5% Physan 20 (Maril Products, Inc., Tustin, CA), and filled them with a sterilized (autoclaved at 80 °C for two 4-h periods with a 48-h resting period in between) 7:3 mixture of Metro-Mix 360 (Sun Gro Horticulture, Bellevue, WA, USA) and all-purpose sand (Quickrete All Purpose Sand, Atlanta, Georgia, USA). I inoculated each mesocosm with a 200 mL layer of live or sterilized field soil (live: n = 4 contemporary environments × 4 microbe histories × 3 field plots/microbe history × 2 mesocosms per field plot = 96 mesocosms; sterile: n = 4 contemporary environments × 8 mesocosms per contemporary environment = 32 mesocosms; total: N = 128 mesocosms). I planted 50 full-sibling families of *C. fasciculata* into these mesocosms. To generate these full-sibling families, seeds were collected from mothers growing in two restored prairies in southwest Michigan (42°28′23′′N, 85°26′50′′W and 42°26′37′′N, 85°18′34′′W) that had been planted with identical prairie seed mixes in 2010 (Shooting Star Native Seeds; Houston County, MN), and controlled crosses were conducted using these seeds in the greenhouse (for details, see Magnoli, 2020). I scarified seeds by nicking the seed coat with a razor blade and imbibed them by submerging in DI water for 3 d, then planted them into inoculated mesocosms. I maintained plants at 30 °C/18 °C for day/night temps on a 15-hr/9-hr L/D cycle, weeded all mesocosms as needed, and watered as needed (generally once daily).

Five weeks after planting, I applied stress treatments similar to those imposed in the field experiment. For the salt stress treatment, I poured 500 mL of a 12 g/L NaCl evenly across each mesocosm. For the herbicide stress treatment, I sprayed ∼800 mL of Roundup (Monsanto, Anvers, Belgium) on plants so that each leaf was nearly dripping. Roundup was applied at a concentration of 0.05 kg active ingredient (glyphosate) per hectare – pilot experiments suggested that this concentration causes visible signs of stress without causing excessive mortality. For the herbivory stress treatment, I used scissors to remove all leaves more than 2.5 cm long.

I measured plant flowering time and specific leaf area (SLA; leaf area/leaf dry mass), which are traits that can respond to both stress and microbes (e.g., Lau & Lennon, 2011; Wagner *et al*., 2014; Panke-Buisse *et al*., 2015). I recorded flowering date for all plants. However, I only measured SLA for plants that were salt-stressed or unstressed, and inoculated with salt, no-stress, or sterile microbe treatments because it was infeasible to measure SLA on all plants. I chose to evaluate SLA in salt stress because it is physiologically similar to drought stress for plants, and plants commonly reduce SLA in response to drought (Wright *et al*., 2001; Ackerly, 2004).

### Statistical analyses

#### Plasticity

All statistical analyses were conducted in R version 4.2.0 (R Core Team, 2022). To test for direct flowering time and specific leaf area (SLA) plasticity in response to stress, I fit linear mixed models that included days to first flower or SLA separately as response variables; contemporary environment (salt, herbicide, herbivory, no stress) as a fixed effect, and plant family and mesocosm as random effects. For tests of direct plasticity, I only included plants inoculated with sterile microbial inoculum (Fig. **1a**) so as not to confound direct plasticity with microbe-mediated plasticity. A significant contemporary environment effect would indicate that plants exhibited direct plasticity (i.e., plant traits shifted in response to stress itself).

To test how live microbial inoculation influences plant flowering time and specific leaf area (SLA) plasticity, I fit linear mixed models that included days to first flower or SLA separately as response variables. I included contemporary environment (salt, herbicide, herbivory, no stress), live inoculation (live, sterile), and their interaction as fixed effects; and plant family, and mesocosm nested in microbial field plot as random effects (I created a dummy variable for field plot in the sterile treatment). For all live vs. sterile comparisons, I compared plants inoculated with sterile inoculant to plants inoculated with microbes from the no-stress control plots (i.e., I did not include microbes from salt-, herbivory-, or herbicide-stressed plots) so as to not confound live-sterile comparisons with microbe history. A significant live inoculation or live inoculation × contemporary environment effect would indicate that plants exhibited microbe-mediated plasticity. Specifically, a significant live inoculation effect would indicate that microbes acted as the environmental factor that elicited a plastic response in plants (i.e., plant traits responded plastically to microbes; Fig. **1b**), while a significant live inoculation × contemporary environment interaction would indicate that microbes altered plant trait responses to the environment (i.e., plant trait plasticity in response to stress was dependent on live inoculation; Fig **1c**). To test for significant differences among stress and microbe treatments, I performed post-hoc tests using the “emmeans” package (Lenth *et al*., 2018).

To test how microbe history influences plant flowering time and SLA plasticity, I fit identical models as above except I included microbe history (salt, herbicide, herbivory, no-stress) as a predictor instead of live inoculation. For these analyses, I excluded the sterile inoculation treatment.

Plasticity is considered adaptive when plastic trait shifts in a particular environment are in the same direction as favored by natural selection in that environment (Jordan, 1991; Nagy, 1997). Therefore, where I detected significant direct or microbe-mediated plasticity, I determined whether these plastic trait shifts were adaptive by comparing the direction of the shift to the direction of natural selection in each environment (e.g., if plants accelerate flowering under stress or when inoculated with a particular microbial community, and earlier flowering is favored by natural selection in that environment, then this plasticity would be considered adaptive). Natural selection was previously estimated in these experimental greenhouse populations (Bolin & Lau, 2024) using standard genotypic selection analyses which involve regressing relative fitness onto standardized trait values (Rausher, 1992).

#### Genetic variation for traits and plasticity

To test how live inoculation affects the expression of genetic variation for plant flowering time, SLA, flowering time plasticity, and SLA plasticity, I fit linear mixed models that included days to first flower or SLA as the response variable; and live inoculation (live vs. sterile), contemporary environment (salt vs. herbicide vs. herbivory vs. no stress for days to first flower; salt vs. no stress for SLA), plant full-sibling family, and all interactions as predictor variables. Greenhouse mesocosm nested in microbial field plot was included as a random effect. A significant live inoculation × family term would indicate that live inoculation influenced the expression of genetic variation for that plant trait, while a significant live inoculation × contemporary environment × family term would indicate that live inoculation influenced the expression of genetic variation for trait plasticity. To test how microbe history affects the expression of genetic variation for plant traits and trait plasticity, I fit identical models except I included microbe history (salt, herbicide, herbivory, no-stress) as a predictor instead of live inoculation.

An increase in genetic variance when plants are inoculated with live or stress-adapted microbes means microbes would accelerate evolutionary response. However, a change in the expression of genetic variation could also be caused by a re-ranking in plant families, rather than a change in variance. For example, if live inoculation alters the expression of genetic variation for flowering time plasticity, families that accelerate flowering when inoculated with live microbes may delay flowering when inoculated with sterile microbes, and vice versa, but with no change in the variance in plasticity among families. Therefore, when live inoculation affected the expression of genetic variation for plant traits, I tested whether these effects were driven by changes in the magnitude of genetic variance vs. a re-ranking of plant families by using Levene’s test for homogeneity of variances in the plant trait (leveneTest function) and by estimating the Spearman’s rank correlation between family mean trait expression in live vs. sterile inoculation (cor.test function) (Heath *et al*., 2020). When live inoculation affected the expression of genetic variation for plant trait plasticity (i.e., significant live inoculation × contemporary environment × family interaction was detected), I used a similar approach except I first calculated plasticity for each family as the ln-response ratio (LRR) = ln[trait expression under stress/trait expression in non-stressful environment]). I then tested for homogeneity of variances and Spearman’s rank correlations in plant plasticity (LRRs) between live vs. sterile microbial treatments. A significant effect of live inoculation on plant trait or plasticity variance would indicate that microbial effects were driven by changes in variance, while a nonsignificant Spearman’s rank correlation would indicate that effects were driven by a re-ranking of plant families. Similar approaches were used to test for effects of microbe history on the expression of genetic variation, except I included microbe history instead of live inoculation in all models.

Models that included all treatments (i.e., all four contemporary environments, and for microbe history models, all four microbe histories) were limited by lower power because only 23 plant families had at least one individual that flowered in each contemporary environment × live inoculation treatment, and only 12 families had at least one individual that flowered in each contemporary environment × microbe history treatment (Table S1). Therefore, I also analyzed control vs. stress environments separately for each contemporary stress. I present results from these simplified models in the main text, and present results from models that include all treatments in the supplement (Tables S2, S3).

#### Genetic correlations for plasticity

To test for genetic correlations between plant plasticity in response to different stressors, and whether live inoculation influenced genetic correlations, I tested for pairwise correlations between plasticity for flowering time in response to each of my three stress treatments when inoculated with a live vs. sterile microbial community. I first quantified flowering time plasticity in response to each stress for each family as the ln-response ratio of flowering time to stress, or ln(days to first flower under stress/days to first flower in non-stressful environments). I then tested whether these family-level ln-response ratios were correlated using the cor.test function. Finally, I tested whether the strength of these genetic correlations differed between live and sterile inoculation using the cocor function in the “cocor” package (Diedenhofen & Musch, 2015). I tested for genetic correlations across stressors in flowering time, but not SLA, because flowering time was the only trait I measured in multiple stressors.

To test for a genetic correlation between phenological and SLA plasticity in response to salt stress (the only stress environment in which I measured both traits), I quantified SLA plasticity in response to salt stress for each family as the ln-response ratio of SLA to stress, or ln(SLA under salt stress/SLA in non-stressful environments). I then tested whether family-level flowering time and SLA ln-response ratios to salt stress were correlated and whether genetic correlations differed between live and sterile inoculation using the same analyses described above.

## RESULTS

### Plasticity

Live microbial inoculation and microbial responses to stress (microbe history) both mediated plant plasticity in response to stress. Herbivory and herbicide stress delayed plant flowering by 6.6 and 6.3 days, respectively, relative to non-stressful environments (pairwise contrasts: herbivory vs. no stress, *P* = 0.001; herbicide vs. no stress, *P* = 0.003; salt vs. no stress, *P* = 0.24), but only when inoculated with a live microbial community, indicating that microbes mediated plant phenological plasticity (direct plasticity, contemp. env.: *P* = 0.38; microbe-mediated plasticity, live inoculation × contemp. env.: *P* = 0.014; Fig. **3a**; Fig. **1c**; Table S4). These microbe-mediated plastic shifts in flowering time were always maladaptive because stress delayed flowering when inoculated with live microbes, but natural selection favored earlier flowering in all four environments (Fig. **4a-c**, green bars) (Bolin & Lau, 2024). However, in non-stressful environments microbe-mediated plasticity was adaptive as plants inoculated with live microbes tended to accelerate flowering in non-stressful environments relative to plants inoculated with a sterile microbial community. By contrast, plants inoculated with live microbes tended to delay flowering under herbicide and herbivory stress relative to plants inoculated with a sterile microbial community, suggesting that microbe-mediated plastic shifts in flowering time were maladaptive in these environments. However, none of these within-contemporary environment contrasts between live and sterile inoculation were significant (pairwise contrasts: no stress, *P* = 0.20; salt, *P* = 0.66; herbicide, *P* = 0.18; herbivory, *P* = 0.44).

**Figure 3.**
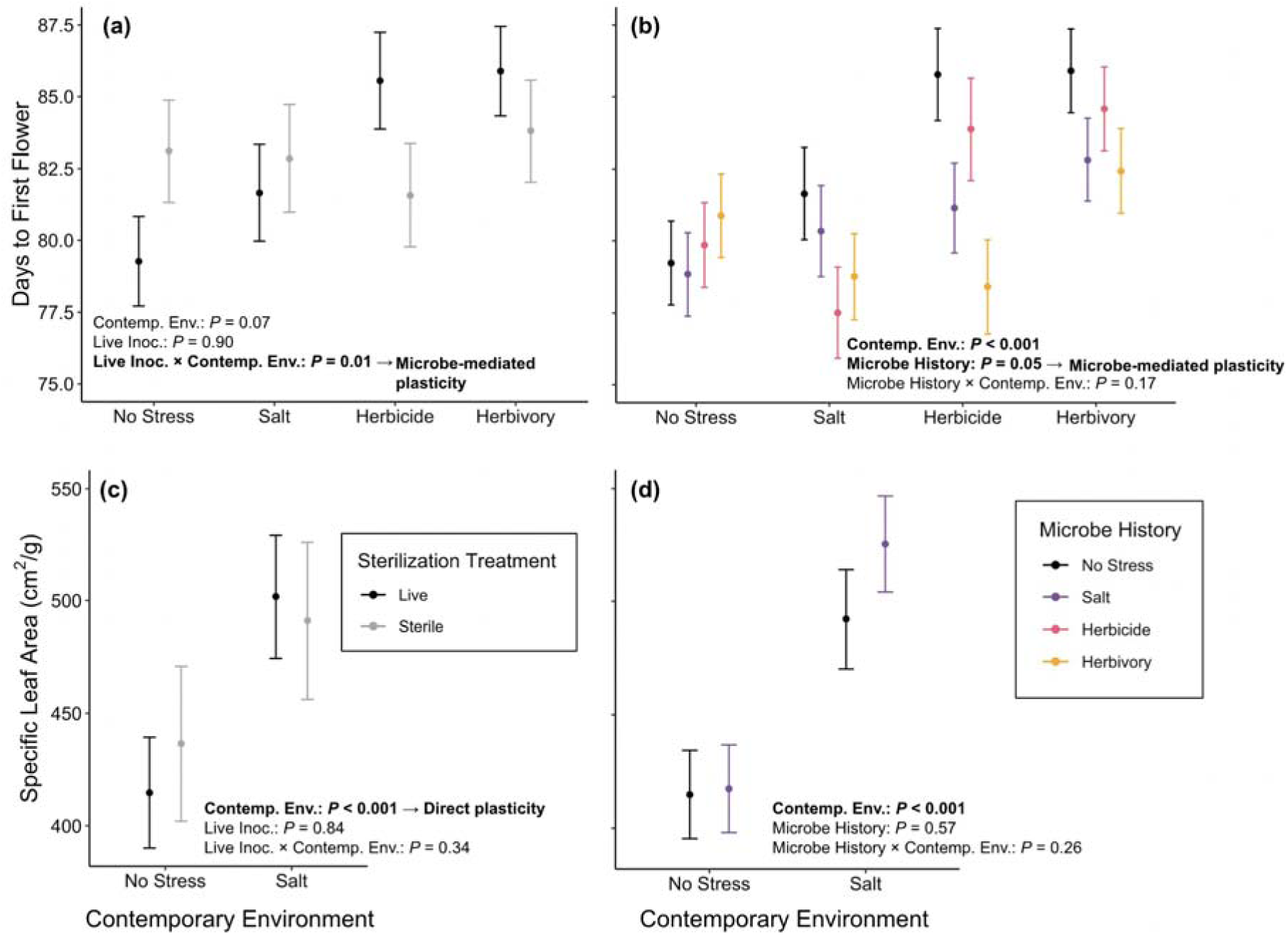
Both live inoculation (“Live Inoc.”) and microbe history mediated plant flowering time plasticity, but not specific leaf area (SLA) plasticity which was only affected by the contemporary environment (“Contemp. Env.”). Plant phenological plasticity was only observed in the presence of a live microbial inoculant (a), and microbial communities that had responded to stress accelerated plant flowering regardless of the contemporary environment (b). In contrast, neither live vs. sterile inoculants nor microbial community responses to stress mediated plant SLA plasticity; SLA plasticity in response to stress was direct (c, d). Estimates and error bars represent estimated marginal means and standard errors. “No stress,” “salt”, “herbicide,” and “herbivory” microbe history treatments are indicated by black, purple, pink, and yellow symbols, respectively, and the sterile treatment is shown in gray.

**Figure 4.**
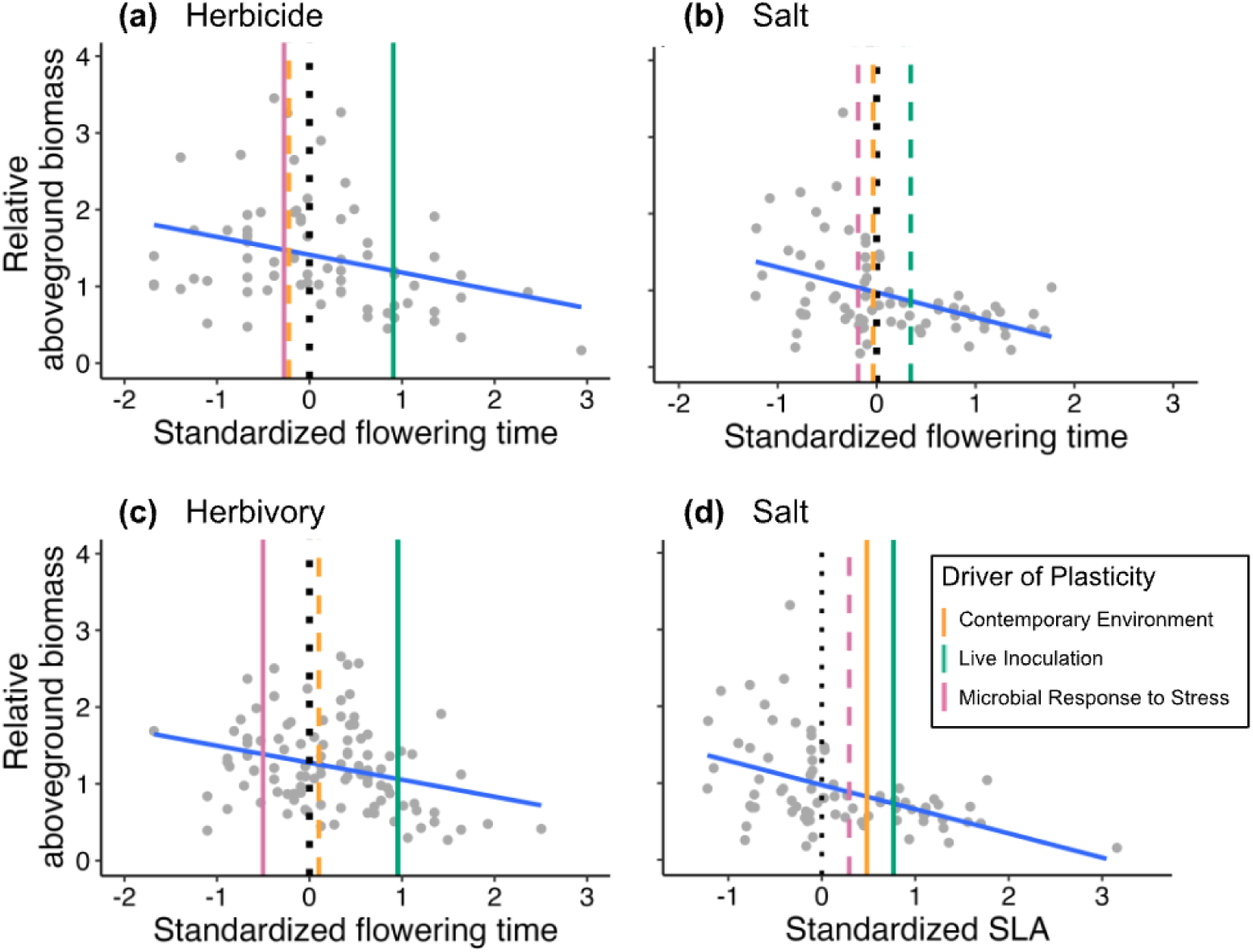
Live inoculation often caused maladaptive trait shifts, while microbial responses to stress often elicited adaptive plasticity. Direct plastic responses to the contemporary environment were much weaker and often maladaptive. Blue lines show the strength and direction of natural selection acting on plant flowering time (a-c) and specific leaf area (SLA; d) under herbicide (a), herbivory (c) and salt stress (b,d). Negative slopes indicate selection for earlier flowering or lower SLA, with steeper slopes indicating stronger selection. Aboveground biomass was relativized by mean aboveground biomass, and traits were standardized by the standard deviation globally (across all treatments). Each point represents a family mean. The dashed vertical line at zero represents the population mean trait value. Plant plasticity driven by the contemporary environment [calculated as (mean trait under stress with sterile inoculation - mean trait under no stress with sterile inoculation)/trait SD; orange], by live inoculation [calculated as (mean trait under stress with live inoculation - mean trait under no stress with live inoculation)/trait SD; green bars], and by microbial responses to stress [calculated as (mean trait under stress with microbes that have responded to stress - mean trait under stress with “no stress” microbes)/trait SD; pink bars] are shown as vertical bars. Solid bars indicate that a driver of plasticity was significant in statistical models, while dashed bars show non-significant trends.

Microbial communities that had responded to stress accelerated flowering relative to soil microbes that had not responded to stress, indicating that stress-adapted microbes mediated plant phenological plasticity by acting as an environment that altered plant trait expression, regardless of the contemporary stress environment (microbe-mediated plasticity, microbe history: *P* = 0.05; Fig. **3c**; Fig. **1b**; Table S5). Herbivory and salt microbes accelerated or tended to accelerate plant flowering by 3 and 2.3 days, respectively (pairwise contrasts: herbivory vs. no stress microbes: *P* = 0.02; salt vs. no stress microbes: *P* = 0.06; herbicide vs. no stress microbes: *P* = 0.16). Thus, while the presence of microbes promoted flowering time shifts that were maladaptive under stress, microbial communities consistently responded to stress in ways that promoted adaptive plant phenological plasticity (Fig. **4a-c**, pink bars).

Contemporary salt stress increased plant SLA by 12%, but this plasticity was unaffected by live inoculation or microbe history (direct plasticity, contemp. env.: *P* = 0.01; Fig. **3b,d**; Tables S6, S7).

Natural selection typically favored lower SLA in both salt stress and non-stressful environments (Bolin & Lau, 2024), making this direct plasticity in response to salt stress maladaptive (Fig. **4d**, black bar).

### Genetic variation for traits and plasticity

Live microbial inoculation and microbe history affected the expression of genetic variation for plant traits (significant live inoculation × family and microbe history × family interactions in Table 1) and trait plasticity (significant live inoculation × contemporary environment × family and microbe history × contemporary environment × family interactions in Table 1) (Figs. S1-S10; Tables S8-S15). In most cases these effects were driven by a re-ranking of plant families in their traits or trait plasticity rather than changes in the amount of genetic variation (i.e., families that delayed flowering under salt stress with sterile inoculation accelerated flowering under stress with live inoculation and vice versa, but there was no change in variance among slopes with live vs. sterile inoculation; Table 1; Fig. S2). However, in one case (flowering time under herbivory stress) microbial effects were driven in part by an increase in variance with live inoculation, and some of this variation with live inoculation was expressed as earlier flowering (the phenotype favored by natural selection), suggesting that a live microbial community may accelerate adaptive phenological evolution in plants (Table 1; Fig. S3).

**Table 1.** Effects of live vs. sterile inoculation and microbe history on the expression of genetic variation for plant trait expression and plasticity.

| Trait | Stress | Microbe Manipulation | Microbes affected expression of genetic variation for plant: | $\chi^2$ from linear mixed models | Driven by | *F* from Levene's Test (significant effect indicates microbes changed variance) | $\rho$ from Spearman's rank correlation ("n.s." indicates microbes caused re-ranking of families) |
| --- | --- | --- | --- | --- | --- | --- | --- |
| Flowering time | Salt | Live vs. Sterile | Trait plasticity | 55 **<sup>a</sup> | Re-ranking of families | 0.09 | 0.05 (n.s.) |
|  |  | Microbe History | Trait expression | 71 *<sup>b</sup> | Re-ranking of families | 0.68 | -0.18 (n.s.) |
|  | Herbicide | Live vs. Sterile | ----- | ----- | ----- | ----- | ----- |
|  |  | Microbe History | Trait plasticity | 31 † | Re-ranking of families | 0.04 | -0.02 (n.s.) |
|  | Herbivory | Live vs. Sterile | Trait expression | 67 * | Change in variance & re-ranking of families | 13 *** | -0.01 (n.s.) |
|  |  | Microbe History | --- | ----- | ----- | ----- | ----- |
| SLA | Salt | Live vs. Sterile | Trait plasticity | 61 * | Re-ranking of families | 0.18 | 0.07 (n.s.) |
|  |  | Microbe History | Trait plasticity | 90 *** | Re-ranking of families | 0.29 | -0.10 (n.s.) |

For rows containing “---”, microbes did not affect the expression of genetic variation for plant trait expression or trait plasticity.
<sup>a</sup> When microbes affected, or tended to affect, the expression of genetic variation for plant trait plasticity, stats are shown for the microbe manipulation (live vs. sterile or microbe history) × contemporary environment × family interaction.
<sup>b</sup> When microbes affected the expression of genetic variation for plant trait expression, stats are shown for the microbe manipulation (live vs. sterile or microbe history) × family interaction.
Full model results are shown in Tables S8-S15.
\*\*\* $p < 0.001$ , \*\* $p < 0.01$ , \* $p < 0.05$ , † $p < 0.1$ .

### Genetic correlations for plasticity

Genetic correlations for phenological plasticity between stressors were common (5 of 6 correlations were statistically significant or marginally significant) and always positive (Table 2), but in one case live inoculation eliminated this genetic correlation (phenological plasticity to herbicide vs. herbivory stress). These positive correlations suggest that plant families that accelerate (or delay) flowering in response to one stress also accelerate (or delay) flowering in response to other stressors.

**Table 2.** Spearman’s correlations between plasticity in response to multiple stressors, and between traits in response to salt stress.

|  |  | Flowering time plasticity in response to: |  |  | SLA plasticity in response to: |
| --- | --- | --- | --- | --- | --- |
|  |  | Herbicide stress | Herbivory stress | Salt stress | Salt stress |
| Flowering time plasticity in response to: | Herbicide stress |  | 0.32 | **0.64\*\*** | — |
|  | Herbivory stress | **0.72\*\*\*** |  | **0.39\*** | — |
|  | Salt stress | 0.38† | **0.37\*** |  | 0.02 |
| SLA plasticity in response to: | Salt stress | — | — | **0.38\*** |  |

Correlations with live and sterile inoculation are shown above and below the diagonal, respectively. Plasticity is calculated as the ln-response ratio to stress, or $\ln(\text{trait under stress}/\text{trait in non-stressful environments})$ . Significant correlations are indicated in bold. † $P < 0.1$ , \* $P < 0.05$ , \*\* $P < 0.01$ , \*\*\* $P < 0.001$ .

Because earlier flowering is advantageous in all these environments (i.e., is favored by natural selection; Bolin & Lau 2024), the disruption of this genetic correlation by microbes would slow the evolution of plasticity in response to simultaneous herbicide and herbivory stress.

I detected a significant positive genetic correlation between phenological and SLA plasticity in response to salt stress, but only with sterile inoculation (live inoculation: *r_14_* = 0.016, *P* = 0.93; sterile inoculation: *r_21_* = 0.38, *P* = 0.027), suggesting that plant families that accelerate flowering under salt stress also decrease SLA under salt stress in the absence of inoculation with a microbial community.

Natural selection favored earlier flowering and lower SLA (Bolin & Lau, 2024), so this correlation would accelerate the evolution of plasticity for both traits in response to salt stress. However, there was no significant difference between live and sterile correlations (comparison of live vs. sterile slopes: Fisher’s *z* = -1.5, *P* = 0.13).

## DISCUSSION

Plant traits are commonly altered by environmental stress, but part of these plant stress responses may in fact be mediated by the diverse soil microbes associating with plants. I found that microbes mediated plant phenological plasticity such that plants only expressed flowering time plasticity when inoculated with a live, non-stress adapted microbial community (Fig. **1c**; Fig. **3a**), while microbial communities that had responded to stress (and therefore are likely to be encountered by plants under stress) accelerated plant flowering across all environments (Fig. **1b**; Fig **3b**). In all cases, microbial community responses to stress promoted adaptive plasticity, pushing plant traits in the direction favored by natural selection in these environments, while direct plasticity in response to the environment (Fig. **1a**) was rarely detected and always maladaptive (Table 3; Fig. 4). My findings also suggest that soil microbes and their responses to stress may influence plant trait evolution by increasing the expression of genetic variation for flowering time and disrupting a genetic correlation for flowering time in response to multiple stressors.

**Table 3.** Summary of microbe-mediated and direct plasticity, and whether plasticity was adaptive (green) or maladaptive (orange).

|  | Flowering Time |  |  | SLA |
| --- | --- | --- | --- | --- |
|  | Salt | Herbicide | Herbivory | Salt |
| Natural selection | **Earlier** | **Earlier** | **Earlier** | **Lower** |
| Direct plasticity | Not Detected | Not Detected | Not Detected | **Higher** |
| Microbe-mediated plasticity (live inoculation) | Not Detected | **Later** | **Later** | Not Detected |
| Microbe-mediated plasticity (microbe history) | Not Detected | **Earlier** | **Earlier** | Not Detected |

Plasticity is considered adaptive when trait shifts are in the direction favored by natural selection. For the natural selection row, each cell denotes the direction of natural selection (Bolin & Lau, 2024). For plasticity rows, each cell denotes the direction of the plastic trait shift between no stress and stress contemporary environments with sterile inoculation (direct plasticity), between no stress and stress contemporary environments with live inoculation (live inoculation-mediated plasticity), and between stress-adapted and non stress-adapted microbes (microbe history-mediated plasticity) detected in this experiment. “Not Detected” indicates that no plasticity of a given types was detected in a given contemporary environment. Adaptive plastic trait shifts are shown in green, maladaptive shifts are shown in orange.

### Microbes mediate plant plasticity

Live inoculation mediated plant flowering time plasticity such that plasticity was only observed when plants were grown in the presence of a live microbial community (Fig. **1c**; Fig. **3a**). Previous studies have shown that specific microbial mutualists such as fungal endophytes, mycorrhizal fungi, and plant growth promoting bacteria can promote (or inhibit) plant plasticity in response to the environment (reviewed in Goh *et al*., 2013), but here I showed that natural diverse microbial communities can also mediate plant plasticity, uncovering a cryptic, but potentially important, role for diverse soil microbes in plant phenotypic responses to the environment. This microbe-mediated phenological plasticity was always maladaptive as plants delayed flowering under stress (which was disfavored by natural selection; Bolin & Lau 2024) when inoculated with a live microbial community. However, microbes that had responded to stress counteracted these maladaptive plastic shifts by accelerating flowering relative to microbial communities that had not responded to stress, suggesting that previously observed patterns of microbe-mediated adaptation (sensu Petipas *et al*., 2021), microbial rescue (sensu Mueller *et al*., 2020), and habitat-adapted symbiosis (sensu Rodriguez *et al*., 2008) may be driven in part by adaptive microbe-mediated plasticity. Specifically, these microbial communities acted as an environmental factor that accelerates plant flowering (Fig. **1b**), counteracting other aspects of the environment that delayed flowering. If these patterns are common, these findings indicate that microbes may be predominant drivers of plant phenological plasticity, and that microbial community responses to stress may commonly protect plants from plastically expressing maladaptive phenotypes under stress.

Microbial responses to stress can mediate plant plasticity in two ways (Fig. **1b,c**), and these likely arise from different biological processes. First, microbes may mediate plant plasticity such that plants only express plasticity in the presence of a live or stress-adapted microbial community (Fig. **1c**, Fig. **3a**) (Goh et al., 2013). In this case, the effect of a microbial community on plant trait expression differs among environments. Thus, this type of microbe-mediated plasticity could arise if microbial trait expression differs among environments (i.e., if microbial trait expression is plastic), or if plant trait expression that affects their interactions with microbes differs among environments (e.g., if plant root traits such as root exudation profiles are plastic; Petipas *et al*. 2020). Furthermore, microbial community composition need not differ among environments, as was likely the case where live inoculation altered plant flowering time (Fig. **3a**), so long as the expression of microbial traits that ultimately alter plant traits (e.g., microbial phytohormone production or traits that alter the soil environment for plants; Friesen *et al*., 2011; Martiny *et al*., 2015) differs among environments. Second, microbes may mediate plant plasticity by acting as an environment that alters plant trait expression regardless of the current environment (Fig. **1b**, Fig. **3b**) (Friesen *et al*., 2011). This could occur if, for example, microbial taxa that become abundant under a certain stress alter plant traits in all environments (again, either directly via phytohormone production or indirectly by altering the soil environment). Here, microbes that had responded to stress accelerated plant flowering in all environments (Fig. **3b**), so stress selected for microbial traits that accelerated flowering, and these microbial traits were not plastic in response to the contemporary environment.

### Microbes alter the evolution of plant plasticity

Soil microbes – both live inoculation and the stress history of the microbial community – affected the expression of genetic variation for plant flowering phenology, indicating that microbes may affect the pace of evolution of flowering time in these populations. In one case, live microbial inoculation exposed cryptic trait variation that was favored by natural selection in the stressful environment (earlier flowering under herbivory stress; Bolin & Lau, 2024), potentially facilitating adaptive evolution (Ghalambor *et al*., 2007). The role of soil microbes in the expression of genetic variation in plants has received little attention (but see O’Brien *et al*., 2019), but my findings suggest that this is another way in which microbes may affect plant evolution, similar to their effects on other evolutionary processes, including plant natural selection and gene flow (reviewed in O’Brien *et al*. 2021). Microbes also often affected the expression of genetic variation for trait plasticity in both plant flowering time and SLA, but these effects were always driven by a re-ranking of plant families in their plastic response to stress rather than in increasing in the magnitude of genetic variation. Therefore, these microbe-mediated changes in the expression of genetic variation are unlikely to affect the response to selection under stress (e.g., Hoffmann & Merilä, 1999).

However, the evolution of plasticity also depends on the strength of genetic correlations (Van Kleunen & Fischer, 2005), and I found that microbes sometimes disrupted (or tended to disrupt) plant genetic correlations for plasticity. I detected significant positive genetic correlations for flowering time plasticity in response to multiple stressors, which would accelerate the evolution of plasticity if natural selection for plasticity is uniform across stressors, or would slow evolution if natural selection for plasticity in response to one stressor opposes selection for plasticity in response to the other stressor.

However, live inoculation disrupted one of these genetic correlation (in response to herbicide and herbivory stress; Table 2), suggesting that microbes may alter the pace of the evolution of flowering time plasticity in response to multiple stressors. Similarly, I detected a positive genetic correlation between flowering time and SLA plasticity in response to salt stress in the absence of a live microbial inoculant, but this genetic correlation disappeared when plants were inoculated with a live microbial community.

While previous studies have investigated genetic correlations in traits underlying plant-microbe interactions (Ossler & Heath, 2018) and in plant drought tolerance between live and sterile microbial treatments (Fitzpatrick *et al*., 2019), my results newly show that microbes can disrupt genetic correlations for plant trait plasticity.

## Implications & Conclusions

Microbes that have responded to stress can alter plant traits, meaning when the environment changes, plant traits are responding to both the environment and correlated changes in the microbial community. If we misattribute the cause of plant plasticity to the environment itself and ignore microbial effects, our predictions about how plants will respond to future stress will likely be inaccurate (Lau & Bolin, 2024). Additionally, plant plastic responses may be unpredictable across space and time due to underlying differences in microbial communities that are exerting their own influences over plant plasticity. While previous studies have shown that stress-adapted microbes can alter plant trait expression in salt (Yuan *et al*., 2019), serpentine (Doubková *et al*., 2012), and drought environments (e.g., Lau & Lennon, 2012; Fitzpatrick *et al*., 2018; Giauque *et al*., 2019; Allsup & Lankau, 2019; Bolin *et al*., 2022; Ricks & Yannarell, 2023) individually, my results show that these microbial community effects may alter plant traits across several stress environments, highlighting the importance of these microbial legacies in determining plant phenotypes more generally.

I showed that microbial communities can respond to several stressors in ways that influence plant traits, but the strength of these effects is likely to depend on both the strength and the duration of the stress event, and how quickly microbial communities respond to that stress event. For example, microbes may not have time to adequately respond to a short or benign stress event in ways that meaningfully affect plant traits. In such cases, live inoculation may matter most for plant phenotypes as differences between stress histories are minimized. But when microbial communities do respond rapidly to stress, as has been observed in less than a week (e.g., Mackelprang *et al*., 2011; Rath *et al*., 2019), plant phenotypic responses to stress may be cryptically driven or offset by their responses to the microbial communities they encounter in these environments.

Overall, I showed that soil microbes and their responses to stress can promote adaptive trait shifts in plants via microbe-mediated plasticity (Fig. **1b,c**). While live microbial inoculation resulted in maladaptive microbe-mediated plasticity in response to stress (Fig. **1c**), microbial communities that had responded to stress reduced these maladaptive plastic shifts, potentially buffering plants from the stress (Fig. **1b**). Finally, microbes altered the expression of genetic variation for plant phenology and disrupted a genetic correlation for plasticity in response to dual stressors. These findings suggest that microbes may cryptically contribute to not only the ecology, but also the evolution of plasticity in plants.

## ACKNOWLEDGMENTS

LGB was supported as an NSF Graduate Research Fellow (GRFP-2141416). I thank Maddie Gellinger, Evan Lacey, Ashley Kovach-Hammons, and Trevor Gress for help with data collection, application of stress treatments, and harvest; John Lemon and Tom Pirtle for greenhouse assistance; Susan Magnoli for seeds; Jennifer Lau, Sarah Fitzpatrick, Jeffrey Conner, and the Lau, Fitzpatrick, and Conner labs for designing and implementing the field experiment from which I sourced soil microbial inocula; and Jennifer Lau, Jay Lennon, Leonie Moyle, Heather Reynolds, Jennifer Rudgers, Emma Boehm, Thomas Zambiasi, and the Lau and Rudgers labs for thoughtful comments on this manuscript.

## SUPPORTING INFORMATION

The following Supporting Information is available for this article:

Table S1 Replication for expression of genetic variation models

Table S2 Factors influencing the expression of genetic variation for plant flowering time across all environments (live inoculation)

Table S3 Factors influencing the expression of genetic variation for plant flowering time across all environments (microbe history)

Table S4 Factors influencing plant flowering time (live inoculation)

Table S5 Factors influencing plant flowering time (microbe history)

Table S6 Factors influencing plant specific leaf area (live inoculation)

Table S7 Factors influencing plant specific leaf area (microbe history)

Table S8 Factors influencing the expression of genetic variation for plant flowering time in salt stress and non-stressful environments (live inoculation)

Table S9 Factors influencing the expression of genetic variation for plant flowering time in herbivory stress and non-stressful environments (live inoculation)

Table S10 Factors influencing the expression of genetic variation for plant flowering time in herbicide stress and non-stressful environments (live inoculation)

Table S11 Factors influencing the expression of genetic variation for plant specific leaf area in salt stress and non-stressful environments (live inoculation)

Table S12 Factors influencing the expression of genetic variation for plant flowering time in salt stress and non-stressful environments (microbe history)

Table S13 Factors influencing the expression of genetic variation for plant flowering time in herbivory stress and non-stressful environments (microbe history)

Table S14 Factors influencing the expression of genetic variation for plant flowering time in herbicide stress and non-stressful environments (microbe history)

Table S15 Factors influencing the expression of genetic variation for plant specific leaf area in salt stress and non-stressful environments (microbe history)

Figure S1 Plots showing the expression of genetic variation for plant flowering time across all environments (live inoculation)

Figure S2 Plots showing the expression of genetic variation for plant flowering time in salt and no-stress environments (live inoculation)

Figure S3 Plots showing the expression of genetic variation for plant flowering time in herbivory and no- stress environments (live inoculation)

Figure S4 Plots showing the expression of genetic variation for plant flowering time in herbicide and no- stress environments (live inoculation)

Figure S5 Plots showing the expression of genetic variation for plant specific leaf area in salt and no- stress environments (live inoculation)

Figure S6 Plots showing the expression of genetic variation for plant flowering time across all environments (microbe history)

Figure S7 Plots showing the expression of genetic variation for plant flowering time in salt and no-stress environments (microbe history)

Figure S8 Plots showing the expression of genetic variation for plant flowering time in herbivory and no- stress environments (microbe history)

Figure S9 Plots showing the expression of genetic variation for plant flowering time in herbicide and no- stress environments (microbe history)

Figure S10 Plots showing the expression of genetic variation for plant specific leaf area in salt and no- stress environments (microbe history)

## REFERENCES

1. Ackerly D. 2004. Functional strategies of chaparral shrubs in relation to seasonal water deficit and disturbance. Ecological Monographs 74: 25–44.

2. Allsup C, Lankau R. 2019. Migration of soil microbes may promote tree seedling tolerance to drying conditions. Ecology 100: e02729.

3. Angulo V, Beriot N, GarciaDHernandez E, Li E. 2022. Plant–microbe ecoLevolutionary dynamics in a changing world. New Phytologist 234: 1919–1928.

4. Beals KK, Moore JAM, Kivlin SN, Bayliss SLJ, Lumibao CY, Moorhead LC, Patel M, Summers JL, Ware IM, Bailey JK et al. 2020. Predicting Plant-Soil Feedback in the Field: Meta-Analysis Reveals That Competition and Environmental Stress Differentially Influence PSF. Frontiers in Ecology and Evolution 8: 191.

5. Bolin LG, Lau JA. 2024. Microbial responses to stress cryptically alter natural selection on plants. New Phytologist 242: 2223–2236.

6. Bolin LG, Lennon JT, Lau JA. 2022. Traits of soil bacteria predict plant responses to soil moisture. Ecology 104: e3893.

7. Brunner I, Herzog C, Dawes MA, Arend M, Sperisen C. 2015. How tree roots respond to drought. Frontiers in Plant Science 6: 547.

8. DeWitt TJ, Scheiner SM. 2004. Phenotypic variation from single genotypes. Phenotypic plasticity: functional and conceptual approaches: 1–9.

9. Diedenhofen B, Musch J. 2015. Cocor: a comprehensive solution for the statistical comparison of correlations. PloS One 10: e0121945.

10. Doubková P, Suda J, Sudová R. 2012. The symbiosis with arbuscular mycorrhizal fungi contributes to plant tolerance to serpentine edaphic stress. Soil Biology & Biochemistry 44: 56–64.

11. Elena SF, Lenski RE. 2003. Microbial genetics: evolution experiments with microorganisms: the dynamics and genetic bases of adaptation. Nature Reviews Genetics 4: 457.

12. Fitzpatrick CR, Copeland J, Wang PW, Guttman DS, Kotanen PM, Johnson MTJ. 2018. Assembly and ecological function of the root microbiome across angiosperm plant species. *Proceedings of the National Academy of Sciences*, USA 115: E1157–E1165.

13. Fitzpatrick CR, Mustafa Z, Viliunas J. 2019. Soil microbes alter plant fitness under competition and drought. Journal of Evolutionary Biology 32: 438–450.

14. Friesen ML, Porter SS, Stark SC, von Wettberg EJ, Sachs JL, Martinez-Romero E. 2011. Microbially Mediated Plant Functional Traits. Annual Review of Ecology, Evolution, and Systematics 42: 23–46.

15. Ghalambor CK, McKAY JK, Carroll SP, Reznick DN. 2007. Adaptive versus non-adaptive phenotypic plasticity and the potential for contemporary adaptation in new environments. Functional Ecology 21: 394–407.

16. Giauque H, Connor EW, Hawkes CV. 2019. Endophyte traits relevant to stress tolerance, resource use and habitat of origin predict effects on host plants. New Phytologist 221: 2239–2249.

17. Goh C-H, Veliz Vallejos DF, Nicotra AB, Mathesius U. 2013. The impact of beneficial plant- associated microbes on plant phenotypic plasticity. Journal of Chemical Ecology 39: 826–839.

18. Graves JL Jr, Tajkarimi M, Cunningham Q, Campbell A, Nonga H, Harrison SH, Barrick JE. 2015. Rapid evolution of silver nanoparticle resistance in Escherichia coli. Frontiers in Genetics 6: 42.

19. Heath KD, Podowski JC, Heniff S, Klinger CR, Burke PV, Weese DJ, Yang WH, Lau JA. 2020. Light availability and rhizobium variation interactively mediate the outcomes of legume-rhizobium symbiosis. American Journal of Botany 107: 229–238.

20. Hoffmann AA, Merilä J. 1999. Heritable variation and evolution under favourable and unfavourable conditions. Trends in Ecology & Evolution 14: 96–101.

21. Jin J, Watt M, Mathesius U. 2012. The autoregulation gene SUNN mediates changes in root organ formation in response to nitrogen through alteration of shoot-to-root auxin transport. Plant Physiology 159: 489–500.

22. Jordan N. 1991. Multivariate analysis of selection in experimental populations derived from hybridization of two ecotypes of the annual plant Diodia teres w. (Rubiaceae). Evolution 45: 1760–1772.

23. Kumar A, Verma JP. 2018. Does plant—Microbe interaction confer stress tolerance in plants: A review? Microbiological Research 207: 41–52.

24. Lau JA, Bolin LG. 2024. The tiny drivers behind plant ecology and evolution. American Journal of Botany 111: e16324.

25. Lau JA, Lennon JT. 2011. Evolutionary ecology of plant-microbe interactions: soil microbial structure alters selection on plant traits. New Phytologist 192: 215–224.

26. Lau JA, Lennon JT. 2012. Rapid responses of soil microorganisms improve plant fitness in novel environments. *Proceedings of the National Academy of Sciences*, USA 109: 14058–14062.

27. Ledón-Rettig CC, Pfennig DW, Crespi EJ. 2010. Diet and hormonal manipulation reveal cryptic genetic variation: implications for the evolution of novel feeding strategies. Proceedings of the Royal Society of London Series B: Biological Sciences 277: 3569–3578.

28. Lenth R, Singmann H, Love J, Buerkner P, Herve M. 2018. Emmeans: Estimated marginal means, aka least-squares means. R package version.

29. Mackelprang R, Waldrop MP, DeAngelis KM, David MM, Chavarria KL, Blazewicz SJ, Rubin EM, Jansson JK. 2011. Metagenomic analysis of a permafrost microbial community reveals a rapid response to thaw. Nature 480: 368–371.

30. Magnoli SM. 2020. Rapid adaptation (or not) in restored plant populations. Evolutionary Applications 13: 2030–2037.

31. Martiny, JBH, Jones SE, Lennon JT, Martiny AC. 2015. Microbiomes in light of traits: A phylogenetic perspective. Science 350: aac9323.

32. Metcalf CJE, Henry LP, Rebolleda-Gómez M, Koskella B. 2019. Why Evolve Reliance on the Microbiome for Timing of Ontogeny? mBio 10: e01496–19.

33. Mueller EA, Wisnoski NI, Peralta AL, Lennon JT. 2020. Microbial rescue effects: How microbiomes can save hosts from extinction. Functional Ecology 34: 2055–2064.

34. Nagy ES. 1997. Selection for native characters in hybrids between two locally adapted plant subspecies. Evolution 51: 1469–1480.

35. O’Brien AM, Sawers RJH, Strauss SY, Ross-Ibarra J. 2019. Adaptive phenotypic divergence in an annual grass differs across biotic contexts. Evolution 73: 2230–2246.

36. O’Brien AM, Ginnan NA, Rebolleda-Gómez M, Wagner MR. 2021. Microbial effects on plant phenology and fitness. American Journal of Botany 108: 1824–1837.

37. Osmond CB, Austin MP, Berry JA, Billings WD, Boyer JS, Dacey JWH, Nobel PS, Smith SD, Winner WE. 1987. Stress physiology and the distribution of plants. *Bioscience*

38. Osmond, C. B., M. P. Austin, J. A. Berry, W. D. Billings, J. S. Boyer, J. W. H. Dacey, P. S. Nobel, et al. 1987. Stress physiology and the distribution of plants. Bioscience 37: 38–48.

39. Ossler JN, Heath KD. 2018. Shared Genes but Not Shared Genetic Variation: Legume Colonization by Two Belowground Symbionts. American Naturalist 191: 395–406.

40. Pandey P, Senthil-Kumar M. 2019. Plant-pathogen interaction in the presence of abiotic stress: What do we know about plant responses? Plant Physiology Reports 24: 541–549.

41. Panke-Buisse K, Poole AC, Goodrich JK, Ley RE, Kao-Kniffin J. 2015. Selection on soil microbiomes reveals reproducible impacts on plant function. The ISME Journal 9: 980–989.

42. Petipas RH, Geber MA, Lau JA. 2021. Microbe-mediated adaptation in plants. Ecology Letters 24: 1302–1317.

43. Queitsch C, Sangster TA, Lindquist S. 2002. Hsp90 as a capacitor of phenotypic variation. Nature 417: 618–624.

44. R Core Team. 2022. *R: A language and environment for statistical computing*. R Foundation for Statistical Computing, Vienna, Austria. http://www.R-project.org/.

45. Rath KM, Maheshwari A, Rousk J. 2019. Linking Microbial Community Structure to Trait Distributions and Functions Using Salinity as an Environmental Filter. mBio 10: e01607–19.

46. Rausher MD. 1992. The measurement of selection on quantitative traits: biases due to environmental covariances between traits and fitness. Evolution 46: 616–626.

47. Ricks KD, Yannarell AC. 2023. Soil moisture incidentally selects for microbes that facilitate locally adaptive plant response. Proceedings of the Royal Society of London Series B: Biological Sciences 290: 20230469.

48. Rodriguez RJ, Henson J, Van Volkenburgh E, Hoy M, Wright L, Beckwith F, Kim Y-O, Redman RS. 2008. Stress tolerance in plants via habitat-adapted symbiosis. The ISME Journal 2: 404–416.

49. Rodriguez R, Redman R. 2008. More than 400 million years of evolution and some plants still can’t make it on their own: plant stress tolerance via fungal symbiosis. Journal of Experimental Botany 59: 1109–1114.

50. Rutherford SL. 2003. Between genotype and phenotype: protein chaperones and evolvability. Nature Reviews Genetics 4: 263–274.

51. Rutherford SL, Lindquist S. 1998. Hsp90 as a capacitor for morphological evolution. Nature 396: 336– 342.

52. Smith SE, Read DJ. 2010. Mycorrhizal Symbiosis. Cambridge, UK: Academic Press.

53. Valladares F, Gianoli E, Gómez JM. 2007. Ecological limits to plant phenotypic plasticity. New Phytologist 176: 749–763.

54. Van Kleunen M, Fischer M. 2005. Constraints on the evolution of adaptive phenotypic plasticity in plants. New Phytologist 166: 49–60.

55. Wagner MR, Lundberg DS, Coleman-Derr D, Tringe SG, Dangl JL, Mitchell-Olds T. 2014. Natural soil microbes alter flowering phenology and the intensity of selection on flowering time in a wild Arabidopsis relative. Ecology Letters 17: 717–726.

56. Wright IJ, Reich PB, Westoby M. 2001. Strategy shifts in leaf physiology, structure and nutrient content between species of high- and low-rainfall and high- and low-nutrient habitats. Functional Ecology 15: 423–434.

57. Xi N, Chu C, Bloor JMG. 2018. Plant drought resistance is mediated by soil microbial community structure and soil-plant feedbacks in a savanna tree species. Environmental and Experimental Botany 155: 695–701.

58. Yuan Y, Brunel C, van Kleunen M, Li J, Jin Z. 2019. Salinity-induced changes in the rhizosphere microbiome improve salt tolerance of Hibiscus hamabo. Plant and Soil 443: 525–537.

