## Supplement for "Soil microbes influence the ecology and evolution of plant plasticity"

Author: Lana G. Bolin

The following Supporting Information is available for this article:

**Table S1** Replication for expression of genetic variation models

**Table S2** Factors influencing the expression of genetic variation for plant flowering time across all environments (live inoculation)

**Table S3** Factors influencing the expression of genetic variation for plant flowering time across all environments (microbe history)

**Table S4** Factors influencing plant flowering time (live inoculation)

**Table S5** Factors influencing plant flowering time (microbe history)

**Table S6** Factors influencing plant specific leaf area (live inoculation)

**Table S7** Factors influencing plant specific leaf area (microbe history)

**Table S8** Factors influencing the expression of genetic variation for plant flowering time in salt stress and non-stressful environments (live inoculation)

**Table S9** Factors influencing the expression of genetic variation for plant flowering time in herbivory stress and non-stressful environments (live inoculation)

**Table S10** Factors influencing the expression of genetic variation for plant flowering time in herbicide stress and non-stressful environments (live inoculation)

**Table S11** Factors influencing the expression of genetic variation for plant specific leaf area in salt stress and non-stressful environments (live inoculation)

**Table S12** Factors influencing the expression of genetic variation for plant flowering time in salt stress and non-stressful environments (microbe history)

**Table S13** Factors influencing the expression of genetic variation for plant flowering time in herbivory stress and non-stressful environments (microbe history)

**Table S14** Factors influencing the expression of genetic variation for plant flowering time in herbicide stress and non-stressful environments (microbe history)

**Table S15** Factors influencing the expression of genetic variation for plant specific leaf area in salt stress and non-stressful environments (microbe history)

**Figure S1** Plots showing the expression of genetic variation for plant flowering time across all environments (live inoculation)

**Figure S2** Plots showing the expression of genetic variation for plant flowering time in salt and no-stress environments (live inoculation)

**Figure S3** Plots showing the expression of genetic variation for plant flowering time in herbivory and no-stress environments (live inoculation)

**Figure S4** Plots showing the expression of genetic variation for plant flowering time in herbicide and no-stress environments (live inoculation)

**Table S1.** Replication for models testing the effects of contemporary environment and microbes (live vs. sterile inoculation and microbe history) on the expression of genetic variation for traits and trait plasticity in each treatment combination. The number of families with at least one individual that flowered or survived to produce leaves is given by “n”.

| **Plant Trait** | **Microbe Comparison** | **Contemp. Env. Comparison** | **n** |
| --- | --- | --- | --- |
| Flowering time | Live, Sterile | Salt, Herbivory, Herbicide, No Stress | 23 |
| Flowering time | Live, Sterile | Salt, No Stress | 34 |
| Flowering time | Live, Sterile | Herbivory, No Stress | 47 |
| Flowering time | Live, Sterile | Herbicide, No Stress | 29 |
| SLA | Live, Sterile | Salt, No Stress | 43 |
| Flowering time | Salt, Herbivory, Herbicide, No Stress | Salt, Herbivory, Herbicide, No Stress | 12 |
| Flowering time | Salt, No Stress | Salt, No Stress | 34 |
| Flowering time | Herbivory, No Stress | Herbivory, No Stress | 44 |
| Flowering time | Herbicide, No Stress | Herbicide, No Stress | 22 |
| SLA | Salt, No Stress | Salt, No Stress | 43 |

**Table S2.** Results of a general linear mixed model testing the effects of contemporary environment (“Contemp. Env.”), live inoculation (“Live Inoc.”), and plant full-sib family (“Family”) on plant flowering time across all four environments.

| **Term** | **df** | ***𝝌^2^*** | ***P*** |
| --- | --- | --- | --- |
| Contemp. Env. | 3 | 7.46 | 0.059 |
| Live Inoc. | 1 | 0 | 0.998 |
| **Family** | **49** | **77.5** | **0.006** |
| **Contemp. Env. × Live Inoc.** | **3** | **11.9** | **0.008** |
| Contemp. Env. × Family | 145 | 161 | 0.179 |
| **Live Inoc. × Family** | **49** | **68.1** | **0.037** |
| Contemp. Env. × Live Inoc. × Family | 110 | 127 | 0.126 |

Notes: Days to first flower was the response variable, and mesocosm nested in field plot from which live soil was sourced were included as random effects (we coded a dummy field plot for the “sterile” treatment). Significant (*P* < 0.05) terms are bolded.

**Table S3.** Results of a general linear mixed model testing the effects of contemporary environment (“Contemp. Env.”), microbe history, and plant full-sib family (“Family”) on plant flowering time across all four environments.

| **Term** | **df** | ***𝝌^2^*** | ***P*** |
| --- | --- | --- | --- |
| **Contemp. Env.** | **3** | **21.6** | **<0.001** |
| Microbe History | 3 | 5.51 | 0.138 |
| **Family** | **49** | **93.8** | **<0.001** |
| Contemp. Env. × Microbe History | 9 | 9.99 | 0.351 |
| Contemp. Env. × Family | 146 | 161 | 0.181 |
| Microbe History × Family | 147 | 146 | 0.519 |
| Contemp. Env. × Microbe History × Family | 340 | 344 | 0.431 |

Notes: Days to first flower was the response variable, and mesocosm nested in field plot from which live soil was sourced were included as random effects. Significant (*P* < 0.05) terms are bolded.

**Table S4.** Results of a general linear mixed model testing the effects of contemporary environment (“Contemp. Env.”) and live inoculation (“Live Inoc.”) on plant flowering time.

| **Term** | **df** | ***𝝌^2^*** | ***P*** |
| --- | --- | --- | --- |
| Contemp. Env. | 3 | 7.15 | 0.067 |
| Live Inoc. | 1 | 0.013 | 0.909 |
| **Contemp. Env. × Live Inoc.** | **3** | **10.5** | **0.014** |

Notes: Days to first flower was the response variable, and plant family and mesocosm nested in field plot from which live soil was sourced were included as random effects (we coded a dummy field plot for the “sterile” treatment). Significant (*P* < 0.05) terms are bolded.

**Table S5.** Results of a general linear mixed model testing the effects of contemporary environment (“Contemp. Env.”) and microbe history on plant flowering time.

| **Term** | **df** | ***𝝌^2^*** | ***P*** |
| --- | --- | --- | --- |
| **Contemp. Env.** | **3** | **24.3** | **< 0.001** |
| **Microbe History** | **3** | **7.78** | **0.051** |
| Contemp. Env. × Microbe History | 9 | 12.9 | 0.166 |

Notes: Days to first flower was the response variable, and plant family and mesocosm nested in field plot from which live soil was sourced were included as random effects (we coded a dummy field plot for the “sterile” treatment). Significant and marginally significant (*P* < 0.1) terms are bolded.

**Table S6.** Results of a general linear mixed model testing the effects of contemporary environment (“Contemp. Env.”) and live inoculation (“Live Inoc.”) on plant specific leaf area (SLA).

| **Term** | **df** | ***𝝌^2^*** | ***P*** |
| --- | --- | --- | --- |
| **Contemp. Env.** | **1** | **16.2** | **< 0.001** |
| Live Inoc. | 1 | 0.040 | 0.841 |
| Contemp. Env. × Live Inoc. | 1 | 0.896 | 0.344 |

Notes: SLA was the response variable, and plant family and mesocosm nested in field plot from which live soil was sourced were included as random effects (we coded a dummy field plot for the “sterile” treatment). Significant (*P* < 0.05) terms are bolded.

**Table S7.** Results of a general linear mixed model testing the effects of contemporary environment (“Contemp. Env.”) and microbe history on plant specific leaf area (SLA).

| **Term** | **df** | ***𝝌^2^*** | ***P*** |
| --- | --- | --- | --- |
| **Contemp. Env.** | **1** | **46.1** | **< 0.001** |
| Microbe History | 1 | 0.329 | 0.566 |
| Contemp. Env. × Microbe History | 1 | 1.25 | 0.264 |

Notes: SLA was the response variable, and plant family and mesocosm nested in field plot from which live soil was sourced were included as random effects (we coded a dummy field plot for the “sterile” treatment). Significant (*P* < 0.05) terms are bolded.

**Table S8.** Results of a general linear mixed model testing the effects of contemporary environment (“Contemp. Env.”), live inoculation (“Live Inoc.”), and plant full-sib family (“Family”) on plant flowering time in salt stress and non-stressful environments.

| **Term** | **df** | ***𝝌^2^*** | ***P*** |
| --- | --- | --- | --- |
| Contemp. Env. | 1 | 0.127 | 0.722 |
| **Live Inoc.** | **1** | **4.861** | **0.027** |
| **Family** | **49** | **72.91** | **0.015** |
| Contemp. Env. × Live Inoc. | 1 | 1.097 | 0.295 |
| Contemp. Env. × Family | 48 | 40.73 | 0.762 |
| Live Inoc. × Family | 47 | 57.05 | 0.150 |
| **Contemp. Env. × Live Inoc. × Family** | **33** | **54.71** | **0.010** |

Notes: Days to first flower was the response variable, and mesocosm nested in field plot from which live soil was sourced were included as random effects (we coded a dummy field plot for the “sterile” treatment). Only the non-stressful and salt stress contemporary environments were included. Significant (*P* < 0.05) terms are bolded.

**Table S9.** Results of a general linear mixed model testing the effects of contemporary environment (“Contemp. Env.”), live inoculation (“Live Inoc.”), and plant full-sib family (“Family”) on plant flowering time in herbivory stress and non-stressful environments.

| **Term** | **df** | ***𝝌^2^*** | ***P*** |
| --- | --- | --- | --- |
| **Contemp. Env.** | **1** | **9.13** | **0.003** |
| Live Inoc. | 1 | 0.0393 | 0.843 |
| Family | 49 | 55.4 | 0.246 |
| **Contemp. Env. × Live Inoc.** | **1** | **7.38** | **0.007** |
| Contemp. Env. × Family | 49 | 45.7 | 0.609 |
| **Live Inoc. × Family** | **49** | **67.4** | **0.042** |
| Contemp. Env. × Live Inoc. × Family | 46 | 53.0 | 0.223 |

Notes: Days to first flower was the response variable, and mesocosm nested in field plot from which live soil was sourced were included as random effects (we coded a dummy field plot for the “sterile” treatment). Only the non-stressful and herbivory stress contemporary environments were included. Significant (*P* < 0.05) terms are bolded.

**Table S10.** Results of a general linear mixed model testing the effects of contemporary environment (“Contemp. Env.”), live inoculation (“Live Inoc.”), and plant full-sib family (“Family”) on plant flowering time in herbicide stress and non-stressful environments.

| **Term** | **df** | ***𝝌^2^*** | ***P*** |
| --- | --- | --- | --- |
| **Contemp. Env.** | **1** | **4.40** | **0.036** |
| Live Inoc. | 1 | 0.147 | 0.702 |
| **Family** | **49** | **82.9** | **0.002** |
| **Contemp. Env. × Live Inoc.** | **1** | **16.5** | **<0.001** |
| Contemp. Env. × Family | 48 | 61.1 | 0.097 |
| Live Inoc. × Family | 48 | 42.8 | 0.685 |
| Contemp. Env. × Live Inoc. × Family | 28 | 32.0 | 0.274 |

Notes: Days to first flower was the response variable, and mesocosm nested in field plot from which live soil was sourced were included as random effects (we coded a dummy field plot for the “sterile” treatment). Only the non-stressful and herbicide stress contemporary environments were included. Significant (*P* < 0.05) terms are bolded.

**Table S11.** Results of a general linear mixed model testing the effects of contemporary environment (“Contemp. Env.”), live inoculation (“Live Inoc.”), and plant full-sib family (“Family”) on plant specific leaf area (SLA) in salt stress and non-stressful environments.

| **Term** | **df** | ***𝝌^2^*** | ***P*** |
| --- | --- | --- | --- |
| **Contemp. Env.** | **1** | **24.0** | **<0.001** |
| Live Inoc. | 1 | **0.0046** | 0.946 |
| Family | 49 | 59.4 | 0.146 |
| Contemp. Env. × Live Inoc. | 1 | 0.643 | 0.423 |
| **Contemp. Env. × Family** | **48** | **72.6** | **0.013** |
| Live Inoc. × Family | 49 | 46.6 | 0.569 |
| **Contemp. Env. × Live Inoc. × Family** | **42** | **60.8** | **0.031** |

Notes: SLA was the response variable, and mesocosm nested in field plot from which live soil was sourced were included as random effects (we coded a dummy field plot for the “sterile” treatment). Only the non-stressful and salt stress contemporary environments were included. Significant (*P* < 0.05) terms are bolded.

**Table S12.** Results of a general linear mixed model testing the effects of contemporary environment (“Contemp. Env.”), microbe history, and plant full-sib family (“Family”) on plant flowering time in salt stress and non-stressful environments.

| **Term** | **df** | ***𝝌^2^*** | ***P*** |
| --- | --- | --- | --- |
| Contemp. Env. | 1 | 1.84 | 0.175 |
| Microbe History | 1 | 0.550 | 0.458 |
| Family | 49 | 57.0 | 0.203 |
| Contemp. Env. × Microbe History | 1 | 0.0369 | 0.848 |
| Contemp. Env. × Family | 46 | 47.371.4 | 0.420 |
| **Microbe History × Family** | **48** | **71.4** | **0.016** |
| Contemp. Env. × Microbe History × Family | 33 | 30.1 | 0.612 |

Notes: Days to first flower was the response variable, and mesocosm nested in field plot from which live soil was sourced were included as random effects. Only the non-stressful and salt stress contemporary environments were included. Significant (*P* < 0.05) terms are bolded.

**Table S13.** Results of a general linear mixed model testing the effects of contemporary environment (“Contemp. Env.”), microbe history, and plant full-sib family (“Family”) on plant flowering time in herbicide stress and non-stressful environments.

| **Term** | **df** | ***𝝌^2^*** | ***P*** |
| --- | --- | --- | --- |
| **Contemp. Env.** | **1** | **23.0** | **<0.001** |
| Microbe History | 1 | 0.0607 | 0.805 |
| **Family** | **48** | **88.3** | **<0.001** |
| Contemp. Env. × Microbe History | 1 | 0.202 | 0.653 |
| Contemp. Env. × Family | 41 | 46.3 | 0.263 |
| Microbe History × Family | 44 | 30.8 | 0.935 |
| Contemp. Env. × Microbe History × Family | 21 | 30.7 | 0.078 |

Notes: Days to first flower was the response variable, and mesocosm nested in field plot from which live soil was sourced were included as random effects. Only the non-stressful and herbicide stress contemporary environments were included. Significant (*P* < 0.05) terms are bolded.

**Table S14.** Results of a general linear mixed model testing the effects of contemporary environment (“Contemp. Env.”), microbe history, and plant full-sib family (“Family”) on plant flowering time in herbivory stress and non-stressful environments.

| **Term** | **df** | ***𝝌^2^*** | ***P*** |
| --- | --- | --- | --- |
| **Contemp. Env.** | **1** | **4.11** | **0.043** |
| Microbe History | 1 | 0.101 | 0.751 |
| **Family** | **49** | **74.7** | **0.010** |
| Contemp. Env. × Microbe History | 1 | 1.56 | 0.212 |
| Contemp. Env. × Family | 49 | 38.0 | 0.871 |
| Microbe History × Family | 49 | 43.0 | 0.712 |
| Contemp. Env. × Microbe History × Family | 43 | 41.0 | 0.557 |

Notes: Days to first flower was the response variable, and mesocosm nested in field plot from which live soil was sourced were included as random effects. Only the non-stressful and herbivory stress contemporary environments were included. Significant (*P* < 0.05) terms are bolded.

**Table S15.** Results of a general linear mixed model testing the effects of contemporary environment (“Contemp. Env.”), microbe history, and plant full-sib family (“Family”) on plant specific leaf area (SLA) in salt stress and non-stressful environments.

| **Term** | **df** | ***𝝌^2^*** | ***P*** |
| --- | --- | --- | --- |
| **Contemp. Env.** | **1** | **54.8** | **<0.001** |
| Microbe History | 1 | 0.400 | 0.527 |
| Family | 49 | 54.8 | 0.265 |
| Contemp. Env. × Microbe History | 1 | 2.31 | 0.128 |
| **Contemp. Env. × Family** | **48** | **80.7** | **0.002** |
| **Microbe History × Family** | **49** | **69.2** | **0.030** |
| **Contemp. Env. × Microbe History × Family** | **42** | **89.5** | **<0.001** |

Notes: SLA was the response variable, and mesocosm nested in field plot from which live soil was sourced were included as random effects. Only the non-stressful and salt stress contemporary environments were included. Significant (*P* < 0.05) terms are bolded.

**Figure S1.** Expression of genetic variation for flowering time plasticity across all four environments with (a) sterile inoculation and (b) live inoculation.

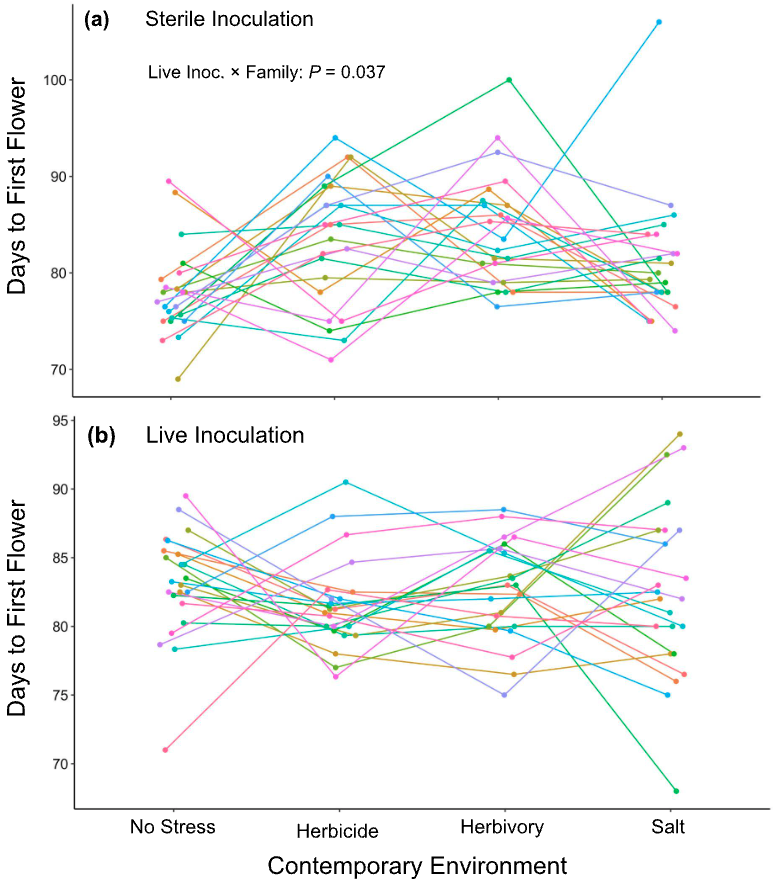

**Figure S2.** Expression of genetic variation for flowering time plasticity in response to salt stress with (a) sterile inoculation and (b) live inoculation.

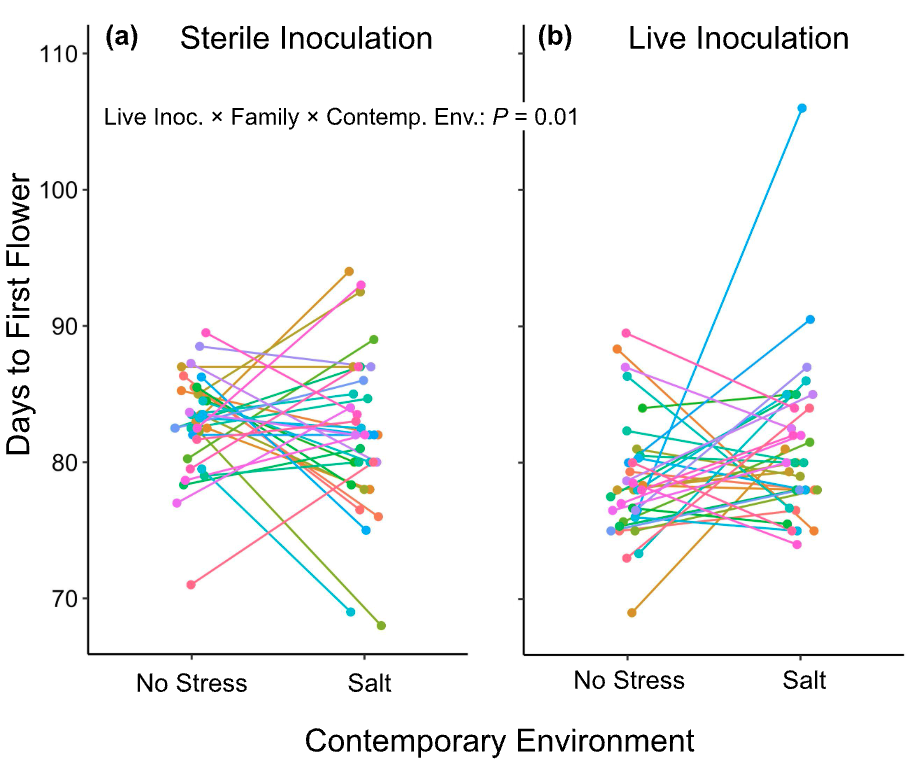

**Figure S3.** Expression of genetic variation for flowering time plasticity in response to herbivory stress with (a) sterile inoculation and (b) live inoculation.

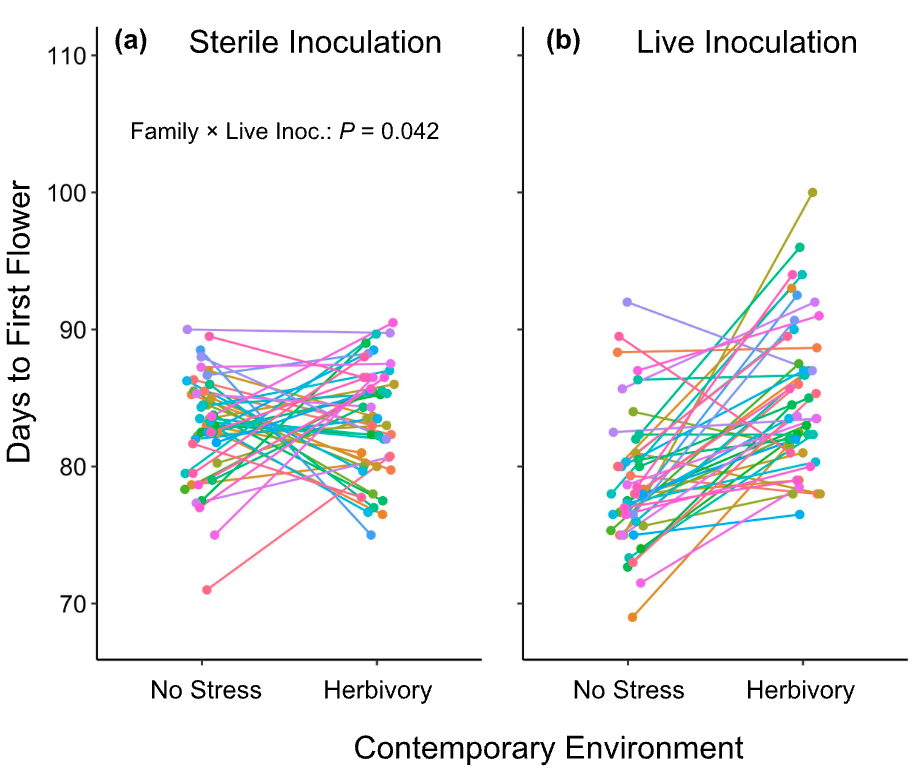

**Figure S4.** Expression of genetic variation for flowering time plasticity in response to herbicide stress with (a) sterile inoculation and (b) live inoculation.

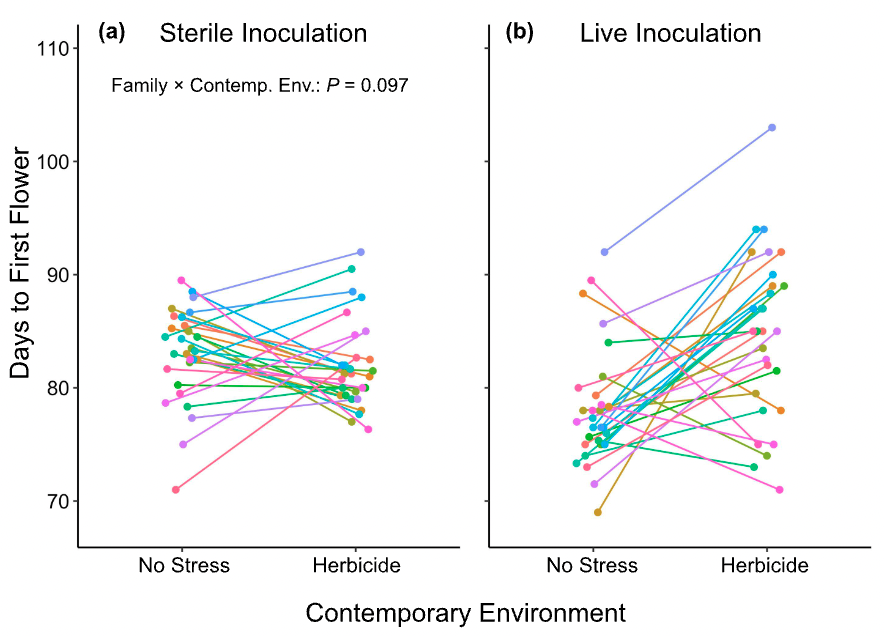

**Figure S5.** Expression of genetic variation for specific leaf area (SLA) plasticity in response to salt stress with (a) sterile inoculation and (b) live inoculation.

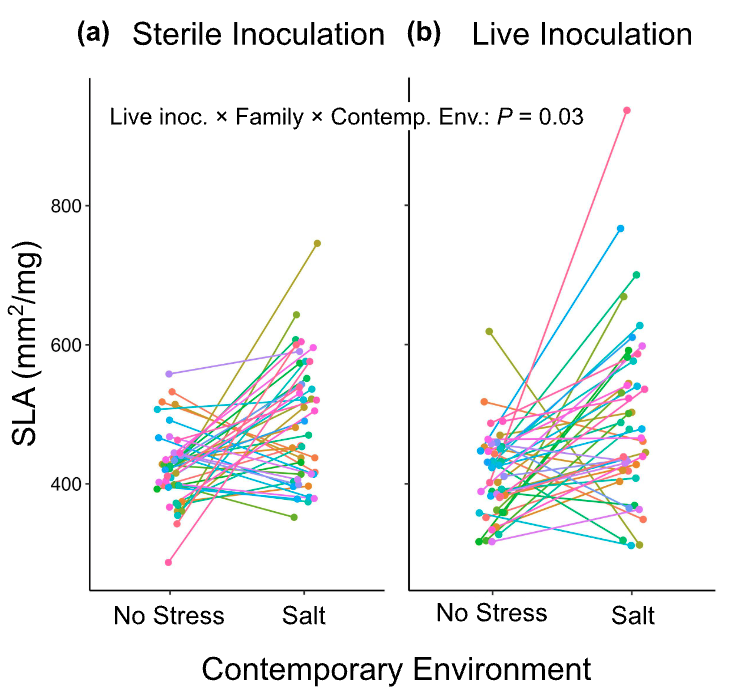

**Figure S6.** Expression of genetic variation for flowering time plasticity across all four environments with microbes from (a) no stress control field plots, (b) microbes from field plots that were treated with salt stress, (c) microbes from field plots that were treated with herbivory stress, and (d) microbes from field plots that were treated with herbicide stress.

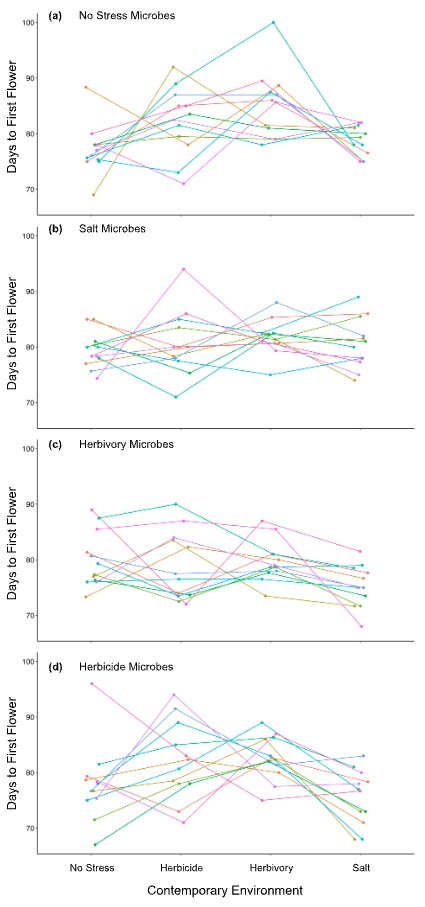

**Figure S7.** Expression of genetic variation for flowering time plasticity in response to salt stress with microbes from (a) no stress control field plots and (b) microbes from field plots that were treated with salt stress.

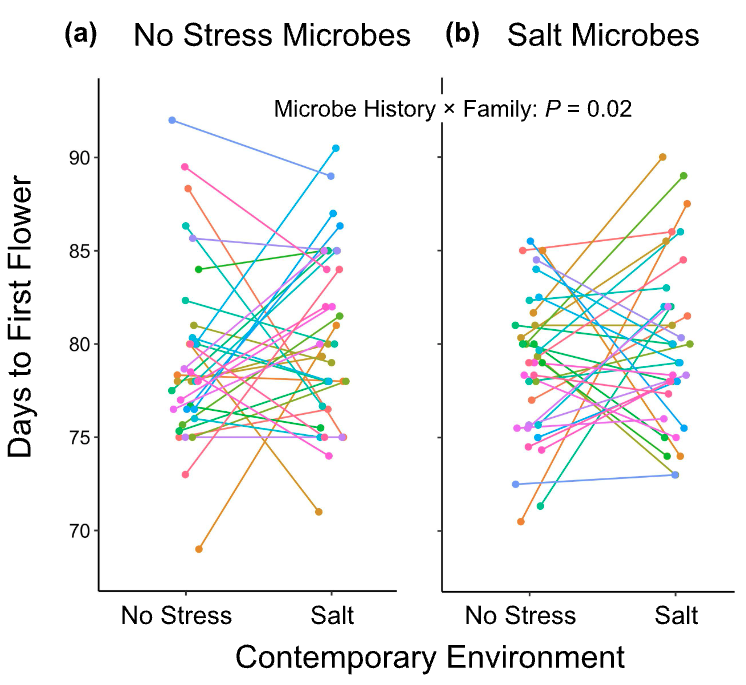

**Figure S8.** Expression of genetic variation for flowering time plasticity in response to herbicide stress with microbes from (a) no stress control field plots and (b) microbes from field plots that were treated with herbicide stress.

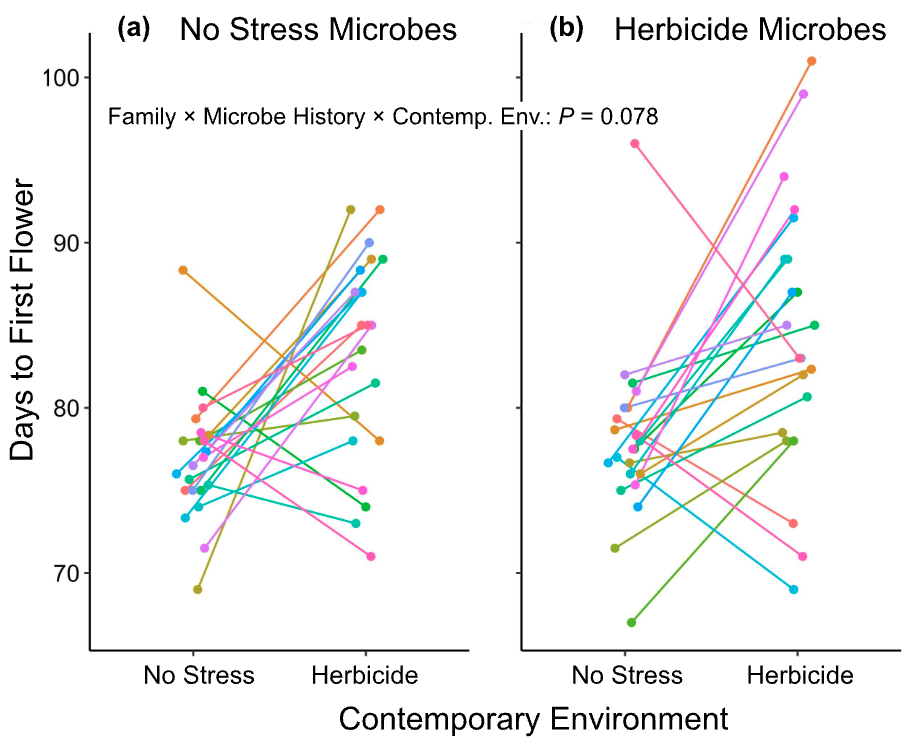

**Figure S9.** Expression of genetic variation for flowering time plasticity in response to herbivory stress with microbes from (a) no stress control field plots and (b) microbes from field plots that were treated with herbivory stress.

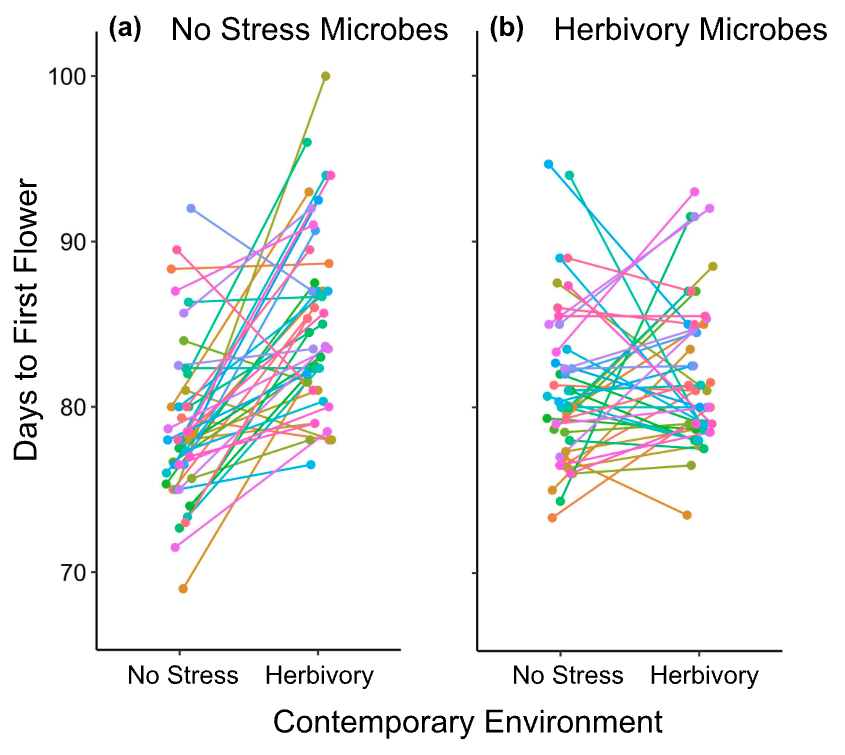

**Figure S10.** Expression of genetic variation for specific leaf area (SLA) plasticity in response to salt stress with microbes from (a) no stress control field plots and (b) microbes from field plots that were treated with salt stress.

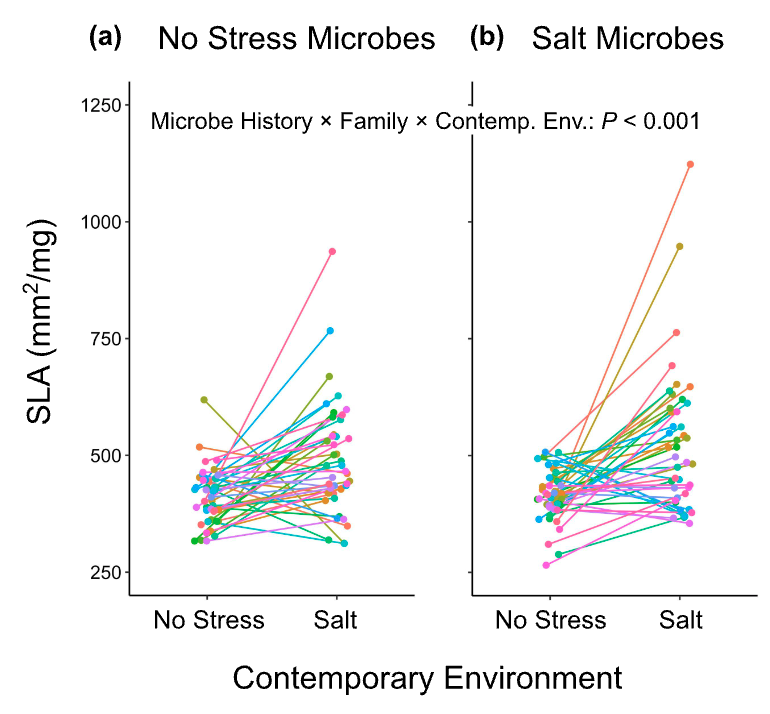
